# Hypoxia-activated fluorescent probes as markers of oxygen levels in plant cells and tissues

**DOI:** 10.1101/2025.02.20.639250

**Authors:** Monica Perri, M. Shahneawz Khan, Antoine L. D. Wallabregue, Viktoriia Voloboeva, Amber M. Ridgway, Edward N. Smith, Hannah Bolland, Ester M. Hammond, Stuart J. Conway, Daan A. Weits, Emily Flashman

## Abstract

- Low oxygen signalling in plants is important in development and stress responses. Measurement of oxygen levels in plant cells and tissues is hampered by a lack of chemical tools with which to reliably detect and quantify endogenous oxygen availability. We have exploited hypoxia-activated fluorescent probes to visualise low oxygen (hypoxia) in plant cells and tissues.
- We applied 4-nitrobenzyl (4NB-) resorufin and methyl-indolequinone (MeIQ-) resorufin to *Arabidopsis thaliana* whole cells and seedlings exposed to hypoxia (1% O_2_) and normoxia (21% O_2_). Confocal microscopy and fluorescence intensity measurements were used to visualise regions of resorufin fluorescence.
- Both probes enter *A.thaliana* whole cells and are activated to fluoresce selectively in hypoxic conditions. Similarly, incubation with *A.thaliana* seedlings resulted in hypoxia-dependent activation of both probes and observation of fluorescence in hypoxic roots and leaf tissue. MeIQ-Resorufin was used to visualise endogenous hypoxia in lateral root primordia of normoxic *A.thaliana* seedlings.
- Oxygen measurement in plants until now has relied on invasive probes or genetic manipulation. Use of these chemical probes to detect applied and endogenous hypoxia has the potential to facilitate a greater understanding of oxygen dynamics in plant cells and tissues, allowing correlation of oxygen concentrations with adaptive and developmental responses to hypoxia.

## Introduction

As aerobic organisms, plants require oxygen for the production of ATP as well as other important biosynthetic processes. While animals have respiratory and circulatory systems for the acquisition and distribution of oxygen, plants rely on photosynthetic oxygen production and diffusion or convection to deliver oxygen to cells and tissues. Regions of low oxygen and oxygen gradients are common in plants (Weits *et al*., 2021) due to high metabolic oxygen consumption, as well as physical barriers to diffusion. The latter arise from hydrophobic and waxy layers present at the surfaces of some tissues which prevent gas diffusion, including in legume root nodules where anaerobic conditions are critical for effective nitrogen fixation (Venado *et al*., 2022), the cuticle and suberized cell layers in seeds in which low oxygen is an important developmental cue (Borisjuk & Rolletschek, 2009; Langer *et al*., 2023) and the deposition of suberin at the surfaces of adventitious roots formed to prevent radial oxygen loss (Shukla & Barberon, 2021). For example, oxygen concentrations of <1 % are reported in seeds (Borisjuk & Rolletschek, 2009), shoot apical meristems and lateral root primordia have metabolically-driven low O_2_ tensions of ∼3 % and <5 % O_2_, respectively (Shukla *et al*., 2019; Weits *et al*., 2019) and O_2_ gradients across *Ricinus communis* (castor bean) stems can range from 21 % at the surface to 7 % in the vascular tissue (van Dongen *et al*., 2003). In addition to endogenous variations in O_2_ tension, plants can be exposed to hypoxia (where oxygen is at a lower concentration than physiologically typical) through pathogen infection (*Botrytis cinerea*-infected Arabidopsis leaves and crown gall tumours arising from infection of wounded tissue by *Agrobacterium tumefaciens* result in O_2_ tensions of 2-5 % (Kerpen *et al*., 2019; Valeri *et al*., 2021)) as well as upon submergence (Bailey-Serres *et al*., 2012).

Low oxygen availability triggers an adaptive response in plants by limiting the activity of oxygen-sensing enzymes, Plant Cysteine Oxidases (PCOs) (Weits *et al*., 2014). This reduces their ability to catalyse oxidation of cysteine residues at the N-termini of their target proteins (White *et al*., 2017), increasing the stability of these proteins by preventing their entering the Cys/Arg branch of the N-degron pathway of protein degradation (Varshavsky, 2019). Primary PCO targets include the Group VII Ethylene Response transcription Factors (ERFVIIs) (Gibbs *et al*., 2011; Licausi *et al*., 2011). When stabilised in low oxygen conditions, the ERFVIIs upregulate the expression of genes which enable acclimation to hypoxia, including fermentative metabolism (Mustroph *et al*., 2009).

Molecular pathways which drive adaptive responses to hypoxia or molecular outcomes of low oxygen niches are well understood, however, the intracellular oxygen concentrations which trigger these responses are not accurately known. To date it has been challenging to accurately quantify oxygen levels in plant cells and tissues non-invasively (Akter *et al*., 2021): the most widely-used method of oxygen measurement is the Clark-type electrode which measures oxygen concentrations at distinct locations (Geigenberger *et al*., 2000; Colmer *et al*., 2020). Mini-electrodes enable resolution to 10 µm, and can reveal cross-sections of O_2_ gradients across tissues (Weits *et al*., 2019), but the electrodes are invasive (thus creating physical damage) and also consume O_2_ locally. Luminescence quenching-based probes have also been used but are similarly invasive, and can be impacted by other quenching agents (Papkovsky, 1993; Pedersen *et al*., 2016; Mori *et al*., 2019; Akter *et al*., 2021). A number of biosensor cassettes have been applied to measure oxygen levels *in vivo* without this invasive treatment (reviewed in (Weits *et al*., 2021)). These have exploited a heterologous mammalian O_2_-sensing system to drive luciferase activity (Iacopino *et al*., 2019) and used a hypoxia responsive promoter element to drive expression of dual fluorescence proteins (Panicucci *et al*., 2020). Although effective, these strategies require significant engineering and are not easily applicable to a range of plant tissues or species. There are no chemical probes reported to date applicable in plants for O_2_ sensing. Since they do not require genetic manipulation or invasive procedures, chemical probes represent a promising system to investigate oxygen-sensing in plant cells and shed light on physiological responses to these conditions. An ideal oxygen-sensing probe should have high sensitivity to hypoxia, potentially finely discriminating among oxygen levels, should be non-toxic, permeable to plant cells and should be detectable in plant systems. Considering that the structure of an elaborate cell wall strictly limits the uptake of large molecules in plants (Tepfer & Taylor, 1981), efficient tissue permeability needs to be tested for chemical probes. In the case of fluorescence-based chemical probes, avoiding florescence spectral overlap between probe and chlorophyll is also a prerequisite consideration for such O_2_-sensing in plants.

Hypoxia-activated imaging agents have been used extensively in mammalian cell and tissue biology to visualise intracellular hypoxia *via* fluorescence or immunostaining (Nordsmark *et al*., 2003; Koch, 2008; Elmes, 2016; Close & Johnston, 2022) . These chemical tools exploit the oxygen-sensitive activity of reductase enzymes: in low oxygen, these enzymes can catalyse bioreduction of redox sensitive functional groups to generate products that can be visualised. One such example is 4-nitrobenzene- resorufin (4NB-Resorufin); hypoxia-dependent bioreduction of 4NB-Resorufin results in cleavage of the 4NB group to release the fluorescent resorufin (Fig. **1**). The proposed mechanism for the bioactivation of resorufin-based probes in mammalian and bacterial systems relies on the fact that the attachment of 4-nitrobenzyl group to the phenol of resorufin results in a non-fluorescent compound. In normoxia, nitroreductase catalyses one electron reduction of the nitro group but the compound is rapidly back oxidized and the resorufin remains uncleaved; however, further reduction of the group occurs under hypoxic conditions leading to the release of the resorufin fluorophore (Collins *et al*., 2018) . Bioreductive moieties such as the nitroaryl groups, quinones, and *N*-oxides have been used in hypoxia activated prodrugs (HAPs) in mammalian systems (McKeown *et al*., 1995; Fryatt *et al*., 1999; Duan *et al*., 2008; Cazares-Korner *et al*., 2013; Hunter *et al*., 2016; Calder *et al*., 2020; Skwarska *et al*., 2021). Additionally, it was shown that isopropyl β-D-1-thiogalactopyranoside (IPTG) in conjugation with the 4-nitrobenzene moiety is inactive, but hypoxia-dependent activation of the molecule releases the IPTG that induces the expression of GFP protein in bacteria (Collins *et al*., 2018).

**Figure 1.**
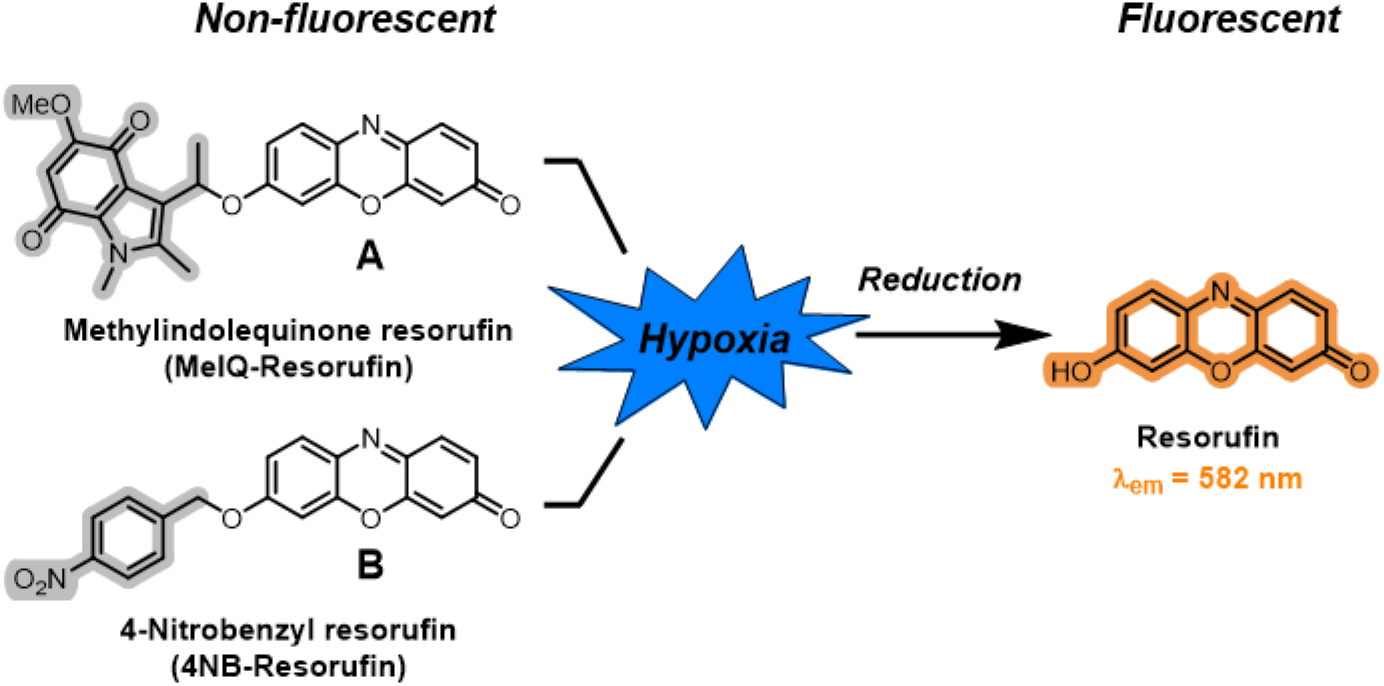
Schematic representation of bioreductive hypoxia activated chemical probes. Methylindolequinone resorufin (MeIQ-Resorufin, **A**) and 4-nitrobenzyl resorufin (4NB-Resorufin, **B**): reduction under hypoxic conditions liberates fluorogenic resorufin.

We considered that hypoxia-activated fluorescent probes could be applied to plant cells and tissues as a way to visualise low oxygen conditions and thus contribute to a clearer understanding of oxygen signalling processes in plants. We investigated two resorufin-based fluorogenic probes, 4NB-Resorufin and a recently reported methyl-indolequinone-conjugated resorufin probe (MeIQ-Resorufin, which has been used in human cancer cell lines (Wallabregue *et al*., 2023)) to determine whether they could permeate plant cells and tissues to report on low oxygen conditions. 4NB-Resorufin has been reported to permeate bacterial cells (Collins *et al*., 2018), therefore we considered the possibility that it might also be able to cross plant cell walls. MeIQ-Resorufin has recently been reported to be activated at higher oxygen concentrations than 4NB-Resorufin (Wallabregue *et al*., 2023) suggesting the possibility that different oxygen concentrations could be discerned in *Arabidopsis thaliana* cell and tissue cultures. Our results reveal that these hypoxia-activated probes are effective at visualising low oxygen conditions (herein termed hypoxia) in this model plant, thus paving the way for an expanded toolkit in detecting and measuring hypoxic conditions in plants.

## Materials and Methods

### MeIQ-Resorufin and 4NB-Resorufin synthesis

MeIQ-Resorufin and 4NB-Resorufin were synthesised as described previously (Wallabregue *et al*., 2023). To yield MeIQ-Resorufin, diisopropyl azodicarboxylate was added dropwise to a solution of 3-(1-hydroxyethyl)-5-methoxy-1,2-dimethyl-1*H*-indole-4,7-dione, resorufin and triphenylphosphine in anhydrous tetrahydrofuran at room temperature under argon. The reaction mixture was stirred vigorously for 24 hours at 50 °C, then concentrated under reduced pressure. The resulting residue was dissolved in ethyl acetate, washed with aqueous sodium hydrogen carbonate, water, aqueous hydrochloric acid (1 M solution), brine, dried (sodium sulfate), filtered, and concentrated under reduced pressure. The crude material was purified using silica gel flash column chromatography. The residue was dissolved in ethyl acetate and precipitated by adding hexane. The precipitation procedure was repeated twice. The precipitate was then dried under reduced pressure to give the title compound as an orange solid.

To prepare 4NB-Resorufin 4-nitrobenzylbromide in *N,N*-dimethylformamide was added to a solution of resorufin and potassium carbonate, stirred at 40 °C for 3 hours, then cooled to room temperature. The solution was diluted with ethyl acetate and washed with brine. The aqueous layer was extracted with ethyl acetate. The organic components were combined, dried (sodium sulfate), filtered and concentrated under reduced pressure. The crude residue was recrystallized from dichloromethane, followed by trituration with diethyl ether to yield the title compound as a red solid. Further details are provided in the Supplementary Information. All the probes were dissolved in DMSO to reach a stock concentration of 1 mM and stored at -20°C, in black Eppendorf tubes to avoid light penetration.

### Plant materials and growth conditions

*A. thaliana* Columbia-0 (Col-0) was used as wild-type ecotype for seedlings. For *in vitro* propagation, sterilized seeds were cultivated on half-strength MS medium (Murashige & Skoog, 1962) supplemented with 1 % (*w/v*) sucrose and 0.8 % (*w/v*) agar, and grown vertically at 22°C, 16:8 day:night photoperiod 100 μmol photons m^-2^ s^-1^ intensity. Two-week old seedlings were used for the experiments. *A. thaliana Landsberg erecta* was used as wild-type ecotype for whole cell cultures; these were propagated in full-strength MS medium, supplemented with 3 % (*w/v*) glucose, 2.5 µg/mL 1- naphthaleneacetic acid (NAA), 0.25 µg/mL kinetin adjusted to pH 5.8, and renewed weekly. Experiments were performed with cells from cultures which had been growing for four days following sub-culture.

### Probe incubation

For incubation with cell cultures, 1 mL of four-day old cell culture was incubated with a final concentration of 10 µM of probe, for two hours (based on studies with similar probes in mammalian cells (Wallabregue *et al*., 2023) as well as time course studies of fluorescence intensity in the presence of each probe, Fig. 3). 1 mL of cell cultures incubated with 1% DMSO and 1 mL of MS incubated with either 1 % DMSO or 10 µM of probe were used as controls. For seedlings treatment, two-week old *A. thaliana* seedlings were incubated with 1 mL of liquid MS medium supplemented with 100 µM of probe unless otherwise mentioned, for six hours, to allow time for tissue entry and diffusion. Seedlings incubated with 10% DMSO, MS incubated with the same amount or probe or DMSO, were used as negative controls. All the experiments were performed using 12-well plates, with three biological replicates per plate, unless otherwise stated. Each biological replicate is represented by 1 mL of Arabidopsis cell suspension or one two-week old seedling for the cell culture and seedling experiments, respectively.

### Hypoxia treatment

All the hypoxic treatments were conducted using a CLARIOstar® plate reader (BMG LabTech) supplemented with an Atmospheric Control Unit (ACU) to regulate O_2_ gas level. Probe sensitivity to hypoxia was evaluated over an O_2_ range from 5 % to 0.2 % using the ACU, flushed with N_2_ (g), for two hours (cell culture treatment) or six hours (seedlings), in the dark; the ACU was turned off for controls in normoxic conditions. Plates were shaken at 100 rpm every 30 seconds before fluorescence detection. For each cycle, 20 readings per well were taken and the mean was calculated. Resorufin was detected upon excitation at 550 nm and emission at 585 nm, using a dichroic filter of 567.5 nm. Two-week old *A. thaliana* Col-0 seedlings expressing a HRPE:nLUC construct (Akter *et al*., 2024) were used for *in vivo* luciferase detection upon treatment under normoxic or hypoxic conditions (1% O_2_ *v/v* O_2_/N_2_), for one hour in the dark, using the CLARIOstar® plate reader (BMG LabTech). Four biological replicates were employed in the experiment and wild-type genotype was used as control. Luminescence was detected using the Nano-Glo® Vivazine™ live cell substrate (Promega), following the manufacturer’s instruction. Measurements were collected for 20 minutes, 7 measurements per minute were taken and the mean was calculated.

### Confocal imaging

Four-day old *A. thaliana* cell cultures and two-weeks old seedlings were used for probe detection after hypoxic treatment (1 % O_2_ *v/v* O_2_/N_2_) for two hours (cell culture) or six hours (seedlings). For propidium iodide (PI) staining, cell cultures were incubated with 10 µg/mL of PI solution for 5 minutes, then mounted on slides. Imaging was performed using either ZEISS LSM 780 (Micron, Department of Biochemistry, University of Oxford) or ZEISS LSM 880 Airyscan microscope (Department of Biology, University of Oxford), equipped with 25 x objective lens, upon laser excitation at 561 nm and collection at 570-590 nm for Resorufin imaging, excitation at 488 nm and collection at 604-622 nm for PI, excitation at 561 nm and collection at 650-750 nm for chlorophyll. For lateral root primordia, *A. thaliana* seedlings were grown on half-strength MS supplemented with 0.5 % sucrose for seven days, then incubated with 100 µM MeIQ-Resorufin for 2 hours in 6-well plates in normoxic (21 % O_2_) or hypoxic (1 % O_2_) conditions. After 10 minutes washing in distilled water, seedlings were imaged using a Zeiss LSM800 Airyscan laser scanning confocal microscope (University of Utrecht) with 561 nm excitation laser light and 565-616 nm emission detection. Fluorescence intensity was calculated using ImageJ software (Schindelin *et al*., 2012). Six measurements were taken for each image, divided between background and signal (three measurements per category). The signal intensity was quantified as the ratio of the mean signal measurement to the mean of the background measurements for each image. This process was repeated for three images per probe, under each experimental condition.

### Root Length and Fresh Weight Measurements

To measure plant tolerance to the probe, *A. thaliana* seedlings were grown vertically on squared plates for seven day and then treated with 100 µM 4NB-Resorufin, MeIQ-Resorufin or resorufin, or 10% DMSO, as negative control, under air and 16:8 photoperiod conditions. After four days of treatment, plates were scanned using EPSON Perfection V750 PRO scanner with a resolution of 720 dots per inch. Primary root length was assessed using ImageJ (Schindelin *et al*., 2012). 21-28 biological replicates were used for root length measurements. Growth rate per day was assessed as the increase in length of the primary root after four days of treatment compared to beginning of treatment. Fresh weight was measured after four days of treatment using four biological replicates, each consisting of seven seedlings.

### Statistical analyses

Statistical analyses were performed using GraphPad Prism 5.04 10.2.3(403) or R Statistical Software (version 4.1.3, Foundation for Statistical Computing, Vienna, Austria). Normal distribution of data was evaluated through the Shapiro-Wilk test. Based on the outcome of the normality test, the appropriate tests were performed to evaluate differences. Additional details are provided in the legend of the correspondent figure.

### Spectroscopy

Fluorescence spectra were recorded using a HORIBA Jobin Yvon FluoroLog3 fluorimeter (Hamamatsu R928 detector and a double-grating emission monochromator). The standard conditions for acquiring emission and excitation spectra are room temperature and steady-stated mode unless otherwise stated. Fluorescence spectra were obtained by using GraphPad Prism software.

UV-visible spectra were recorded on a V-770 UV-Visible/NIR Spectrophotometer equipped with Peltier temperature controller and stirrer using disposable polystyrene cuvettes of 1 cm path length. Experiments were conducted at 25 °C unless otherwise stated. UV-visible spectra were plotted using GraphPad software.

pH measurements of solutions were taken using an Oakton pH meter.

### Reactive oxygen species preparation and assays

#### General procedure for hydrogen peroxide (H_2_O_2_) assay

To a quartz cuvette on ice (0–4 °C) containing Milli-Q water and H₂O₂ solution, a probe solution was added. The cuvette was sealed and left at room temperature for 5 minutes. At this point, the final concentrations of H₂O₂ and the hypoxia-imaging probe were 100 µM and 1 µM, respectively. The fluorescence spectra were then measured using the specified parameters at the given time point. Further details are provided in the Supplementary Information.

#### General procedure for hydroxyl radical (HO•) assay

##### Preparation

The hydroxyl radical (HO•) was generated using the Fenton reaction. Ferrous chloride (1 eq.) was added to a solution of H₂O₂ (10 eq.). Upon mixing, the solution turned orange and was used immediately for analysis.

##### Assay

To a quartz cuvette on ice (0 – 4 °C) containing Milli-Q water and generated HO• solution, probe solution was added. At this stage, the final concentrations of HO• and the hypoxia-imaging probe were 100 µM and 1 µM, respectively. The solution was homogenized by pipetting for 10 seconds, then the cuvette was sealed and left at room temperature for 5 minutes. Fluorescence spectra were measured under the specified parameters. Further details are provided in the Supplementary Information.

#### General procedure for light stability assay

To a vial containing DMSO, a probe solution was added. An aliquot was taken from the reaction solution and transferred to a quartz cuvette, followed by the addition of Milli-Q water. The solution was homogenized by pipetting, then the fluorescence spectra were recorded using the specified parameters, serving as the initial time point (T = 0, where T represents time).

The vial was then sealed and placed in an aluminum box, positioned 20 cm away from a white LED and illuminated for 4 hours. After illumination, another aliquot was collected, transferred to a quartz cuvette, and diluted with Milli-Q water. The fluorescence spectrum was subsequently recorded under the specified parameters. Further details are provided in the Supplementary Information.

#### General procedure for the assay with high ionic strength buffers

To a quartz cuvette containing buffer with the desired NaCl concentration, a probe solution was added. The solution was homogenized by pipetting for 10 seconds, then the cuvette was sealed and left at room temperature for 10 minutes. The fluorescence spectra were then recorded using the specified parameters. Further details are provided in the Supplementary Information.

### General procedure for the assay with different pH buffers

To a quartz cuvette containing buffer at the desired pH, a probe solution was added. The solution was homogenized by pipetting for 10 seconds, then the cuvette was sealed and incubated at room temperature for 5 or 60 minutes. The fluorescence spectra were then recorded using the specified parameters. Further details are provided in the Supplementary Information.

## Results

### 4NB-Resorufin and MeIQ Resorufin can penetrate Arabidopsis thaliana whole cells and show hypoxia-dependent activation

We first set out to determine whether the 4NB- and MeIQ-Resorufin probes could permeate *Arabidopsis thaliana* cells. Whole autotrophic suspension-cultured cells were grown for 4 days at 22 °C then incubated with either 10 µM probe or 1 % DMSO loading control prior to shaking in the dark for 2 hours under normoxia (21 % O_2_) or hypoxia (1 % O_2_). Cells were visualised with an inverted scanning confocal microscope with laser excitation at 561 nm and emission detection at 570-590 nm. Chlorophyll fluorescence was measured to facilitate intracellular localisation of resorufin fluorescence and was detected upon excitation at 561 nm and detection at 650-750 nm. Propidium iodide (PI) staining was exploited for live/dead cell imaging, since this dye is not cell-permeable in intact membranes (Riccardi & Nicoletti, 2006); PI signal was detected upon excitation at 488 nm and emission detection at 604-622 nm.

The arising images (Fig. **2a**, Fig **S1**) and signal quantification (Fig, **2b**) show that when cells were incubated with either 4NB-Resorufin or MeIQ-Resorufin in hypoxic conditions, significant increase in fluorescence in the 570-590 nm range was observed in cytosolic regions of the cell, consistent with resorufin fluorescence (Fig. **2b, c**). No fluorescence was observed at this wavelength range when cells were incubated with either 4NB-Resorufin or MeIQ-Resorufin in normoxia (Fig. **2a**). Similarly, no fluorescence was observed if cells were incubated in hypoxia and treated with 1 % DMSO only, nor if either probe was incubated in MS media, in hypoxia and in the absence of cells (Fig. **S2**). Chlorophyll autofluorescence was also observed in the presence of both 4NB-Resorufin and MeIQ-Resorufin and when cells had been incubated in both normoxia and hypoxia. PI fluorescence was used to demonstrate that the probes were entering cells and that the cells remained viable. PI fluorescence emission was only observed at cell surfaces, indicating the probes were not toxic to the cell under the conditions tested. PI staining was then used to determine the proportion of dead or damaged cells as a function of increasing concentrations of each probe (Fig. **S3a, b**). Across the different concentrations tested, there was no significant increase in PI staining with increasing concentration of probe, relative to DMSO control. There was some increase in PI staining in hypoxic control cells compared to normoxic control cells. Overall, the results indicate that, under the conditions used, the presence of the probes does not cause significant damage to the cell membranes and is no more damaging to cells than hypoxia itself.

**Figure 2.**
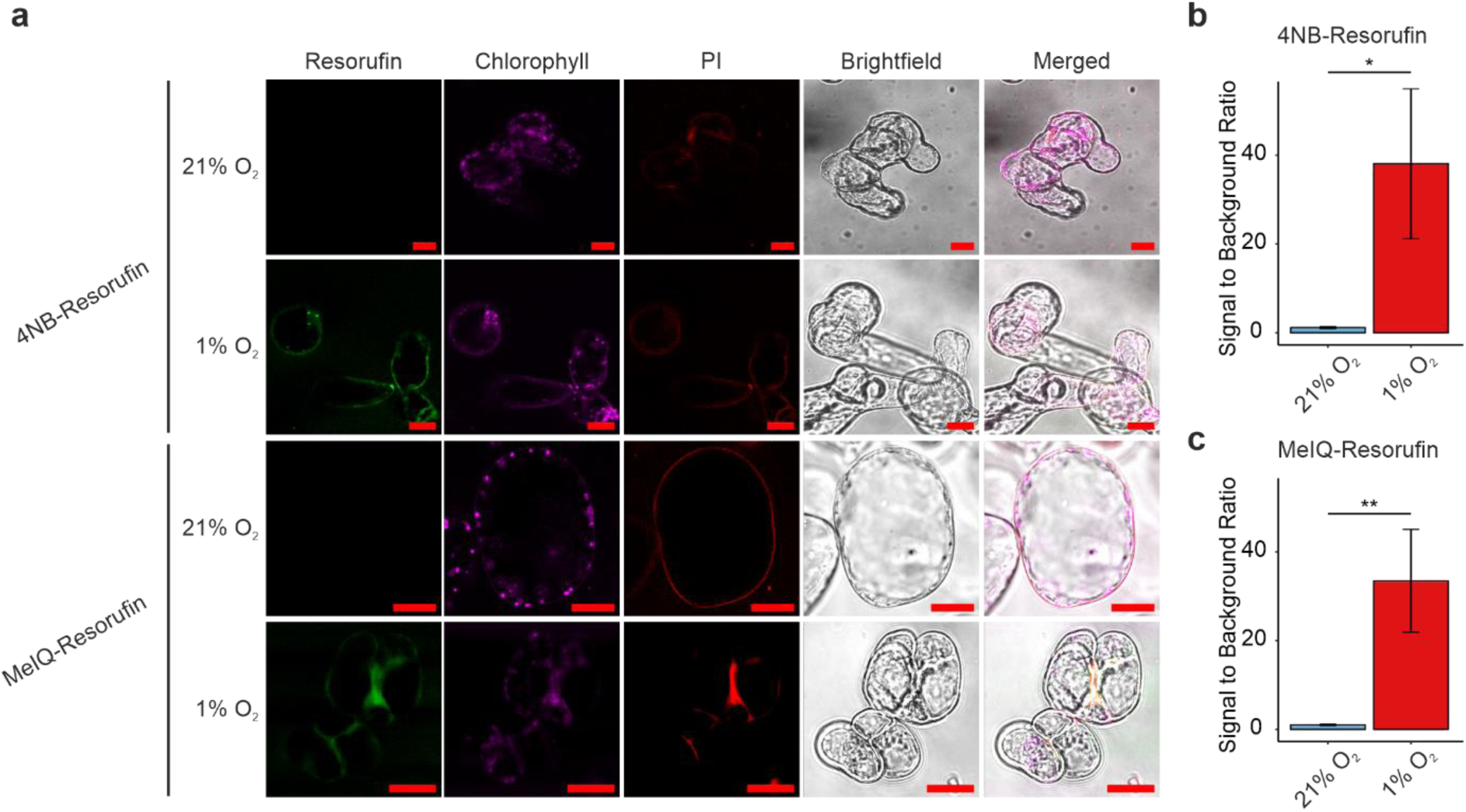
Hypoxic incubation of *Arabidopsis thaliana* cells with 4NB-Resorufin and MeIQ-Resorufin results in fluorescence. (**a**) 1 mL *Arabidopsis thaliana* whole cells were incubated with 10 µM 4-NB-resorufin or MeIQ-Resorufin at 21 % or 1 % O_2_ for 2 hours. Cells were examined for fluorescence of resorufin (green, 570-590 nm), chlorophyll (purple, 650-750 nm) and propidium iodide (red, 604-622 nm). Images show representative cells from ≥ 3 x wells containing 1 mL cells; scale bars: 20 µm. (**b, c**) Quantification of resorufin fluorescence intensity in cells treated with 4NB- or MeIQ-Resorufin upon hypoxia or normoxia represented by the ratio of the signal intensity to the background (*n* ≥ 3). Statistical significance was determined using two-sided Student’s t-test (* p-value < 0.05, ** p-value < 0.01).

The results therefore indicate that both probes are able to permeate Arabidopsis whole cells and enter the cytoplasm. The observed fluorescence in hypoxia but not in normoxia indicates that the probes have been bioreductively cleaved to form resorufin in a manner that is dependent on a sustained decrease in oxygen availability. This is consistent with the bioreductive properties of these probes in bacterial and mammalian systems (Collins *et al*., 2018; Wallabregue *et al*., 2023). Although the reductase enzyme(s) responsible for bioreduction of the probes have not been identified in plants, the same probes are reduced by nitroreductase and CYP450 enzymes in mammalian and bacterial cells (O’Connor *et al*., 2016; O’Connor *et al*., 2017; Collins *et al*., 2018; Wallabregue *et al*., 2023), suggesting that similar enzymes may be enabling probe activation in plant cells. These probes are therefore able to detect whether whole plant cells grown in culture are hypoxic.

### Quantitative fluorescence detection of hypoxia-dependent bioreduction of 4NB-Resorufin and MeIQ-Resorufin in a plate-based format

We next wanted to determine whether the probes could be exploited to detect cellular hypoxia in a higher-throughput plate-based format by measuring fluorescence intensity, and whether such an assay can be used to quantify fluorescence levels as a function of O_2_ concentration. By exposing cells to different concentrations of O_2_ we also wanted to determine whether the bioreduction and probe fluorescence were the same for both probes. 1 mL of autotrophic *A. thaliana* four-day old cell cultures were transferred to 12-well plates and incubated for 2 hours in a plate reader where the atmospheric environment was controlled at concentrations of O_2_ ranging from 0.2 % to normoxia (21 % O_2_). Plates were kept in dark conditions throughout the experiment to avoid O_2_ production through photosynthesis, therefore glucose was added to the culture medium to maintain cells in a heterotrophic state. Fluorescence intensity was measured over time for each condition (Fig. **3a, b**) and average intensity (from 15-120 minutes) was used to determine overall fluorescence levels (Fig. **3c, d**) at each O_2_ concentration.

**Figure 3.**
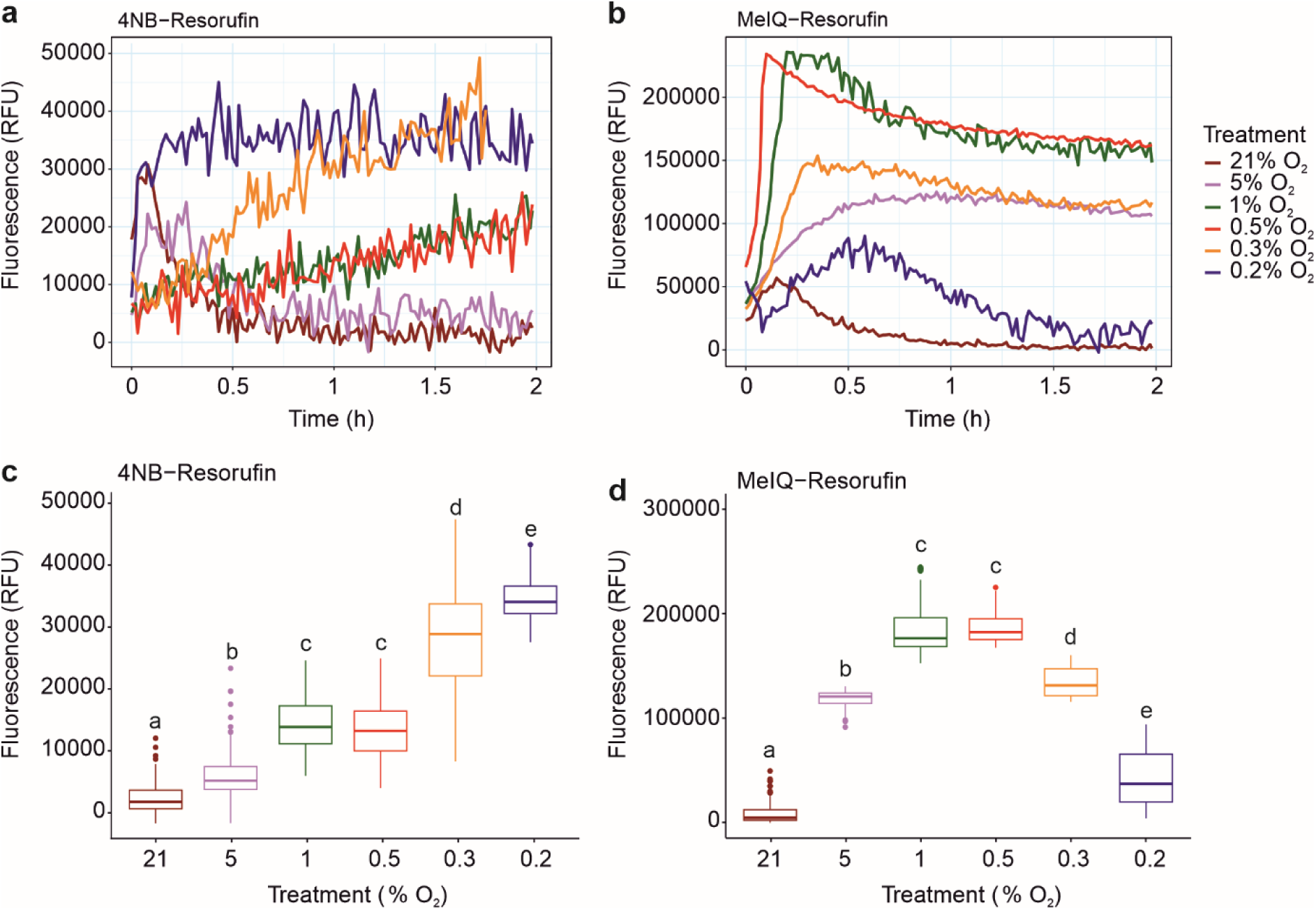
Fluorescence intensity of bioreduced probes varies with O_2_ concentration. Hypoxic incubation of *Arabidopsis thaliana* cells with 4NB-Resorufin (**a**) or MeIQ-Resorufin (**b**) shows variation in O_2_-dependent activation of fluorescence with time. Averaged fluorescence intensity (across the duration of 15-120 minutes) shows a difference in the O_2_ concentration at which the most intense signal is achieved for 4NB-Resorufin (**c**) and MeIQ-Resorufin (**d**). In all experiments, 1 mL *Arabidopsis thaliana* whole cells were incubated with 10 µM of the probes at varying O_2_ concentrations. Data in boxplots (**c**) and (**d**) were taken from ≥3 independent experiments; one-way ANOVA analysis followed by Tukey HSD *post-hoc* test revealed that fluorescence intensity was significantly different from that measured in normoxia in all cases (letters indicate statistical differences between groups, p-value < 0.001).

As detected using microscopy, resorufin fluorescence was observed when cells were equilibrated with both probes in hypoxia compared to cells exposed to normoxia. Cells incubated with 4NB-Resorufin at O_2_ concentrations of 1 % and below showed a sustained increase in fluorescence over the course of 2 hours; when the O_2_ concentration was 0.2 %, a maximum and sustained level of fluorescence was observed after approximately 15 minutes (Fig. **3a**). There was an overall increase in fluorescence intensity as O_2_ concentration reduced from 1 % to 0.2 % (Fig. **3c**). When cells were incubated with MeIQ-Resorufin, a more rapid increase in fluorescence was observed at all O_2_ concentrations measured except 21 % (Fig. **3b**). The absolute levels of fluorescence attained appeared to be dependent on O_2_ concentration, with highest levels of fluorescence observed for 1 % and 0.5 % O_2_ (Fig. **3d**). This fluorescence was not sustained, however, and over the course of 2 hours, fluorescence levels decreased, most notably in the presence of 0.2% O_2_.

MeIQ-Resorufin derived fluorescence appeared to be detected at higher O_2_ concentrations compared to fluorescence observed in the presence of 4NB-Resorufin at equivalent O_2_ levels (Fig. **3b, d**), consistent with comparison of these probes in mammalian cells (Wallabregue *et al*., 2023), suggesting MeIQ-Resorufin may be capable of detecting milder hypoxia than 4NB-Resorufin. However, absolute levels of fluorescence were approximately 5-fold higher in the presence of MeIQ-Resorufin compared to 4NB-Resorufin, which may reflect greater uptake of this probe into the cell, faster intracellular diffusion or a greater propensity for MeIQ-Resorufin bioreduction in the *A. thaliana* cell environment. In line with this, the observed decline in fluorescence signal with time for MeIQ-Resorufin suggests it is sequestered more rapidly: resorufin fluorescence is quenched by acidic conditions (pH 6.5 and below) (Fig. **S4**) suggesting the decline in fluorescence could be a result of vacuolar sequestration of high levels of resorufin derived from the MeIQ-conjugated probe. Alternatively, resorufin fluorescence quenching could derive from reduced intracellular pH associated with lactate fermentation under hypoxia (Wagner *et al*., 2019). Overall, the results indicate that the probes are capable of detecting intra-cellular hypoxia in a semi-quantitative manner in a plate-based assay, thus extending their utility as hypoxia-sensing probes.

### 4NB-Resorufin and MeIQ-Resorufin can detect hypoxic conditions in Arabidopsis thaliana seedling tissues

To confirm the biological relevance of 4NB- and MeIQ-Resorufin as hypoxia probes, we next sought to determine whether cell permeability and bioreduction of the probes could be observed in *A. thaliana* whole seedlings under hypoxic conditions. We anticipated that additional incubation time would be required for diffusion and uptake of probe in seedlings compared to cells; given we also observed a decrease in resorufin fluorescence in whole cells after prolonged incubation with MeIQ-Resorufin at low O2 concentrations, we first determined whether a stable fluorescence signal could be observed in seedlings incubated with 4NB- and MeIQ-Resorufin at 0.1 % O2 over the course of six hours. The fluorescent signal intensity was not strong in cultured cells (Fig. **2**), therefore, to ensure the detection of signal in a more complex system, we increased the probe concentration in these experiments to 100 µM. The arising data (Fig. **S5**) revealed a time-dependent increase in resorufin fluorescence intensity over 6 hours, suggesting that the probes were sufficiently stable to elicit a detectable signal in the whole-seedling environment.

We therefore proceeded to examine seedling tissues exposed to normoxia (21% O2) or hypoxia (1 % O2) by confocal microscopy following incubation with both probes (Fig. **4**). When seedlings were incubated with 4NB-Resorufin (Fig. **4(I)**), hypoxia-dependent fluorescence was observed in root tissue (Fig. **4(I)****a, b**) and leaves (Fig. **4(I)****c, d**). In the root tissue, fluorescence was observed in the cortex and vascular tissue; in the leaf tissue, fluorescence was observed, albeit not homogenously. Overall, these images indicate that 4NB-Resorufin was able to permeate the seedling tissue and translocate through the plant, likely *via* the vasculature. MeIQ-Resorufin was similarly able to permeate seedling tissue and fluoresce in a hypoxia-dependent manner. MeIQ-Resorufin derived fluorescence was observed in vascular regions of the rootand near-homogenously in the leaf tissue (Fig. **4****(II) a-d**). These data suggest translocation of the probe throughout the plant tissue including to the leaf mesophyll layer. As observed in cells, a significant increase in fluorescence was only observed in the presence of either probe (and hypoxia) (Fig. **4e, f**); seedlings treated with equivalent volumes of DMSO did not demonstrate fluorescence at 570-590 nm in either hypoxia or normoxia, while chlorophyll autofluorescence was observed in these seedlings in leaf tissue (Fig. **S6**).

**Figure 4.**
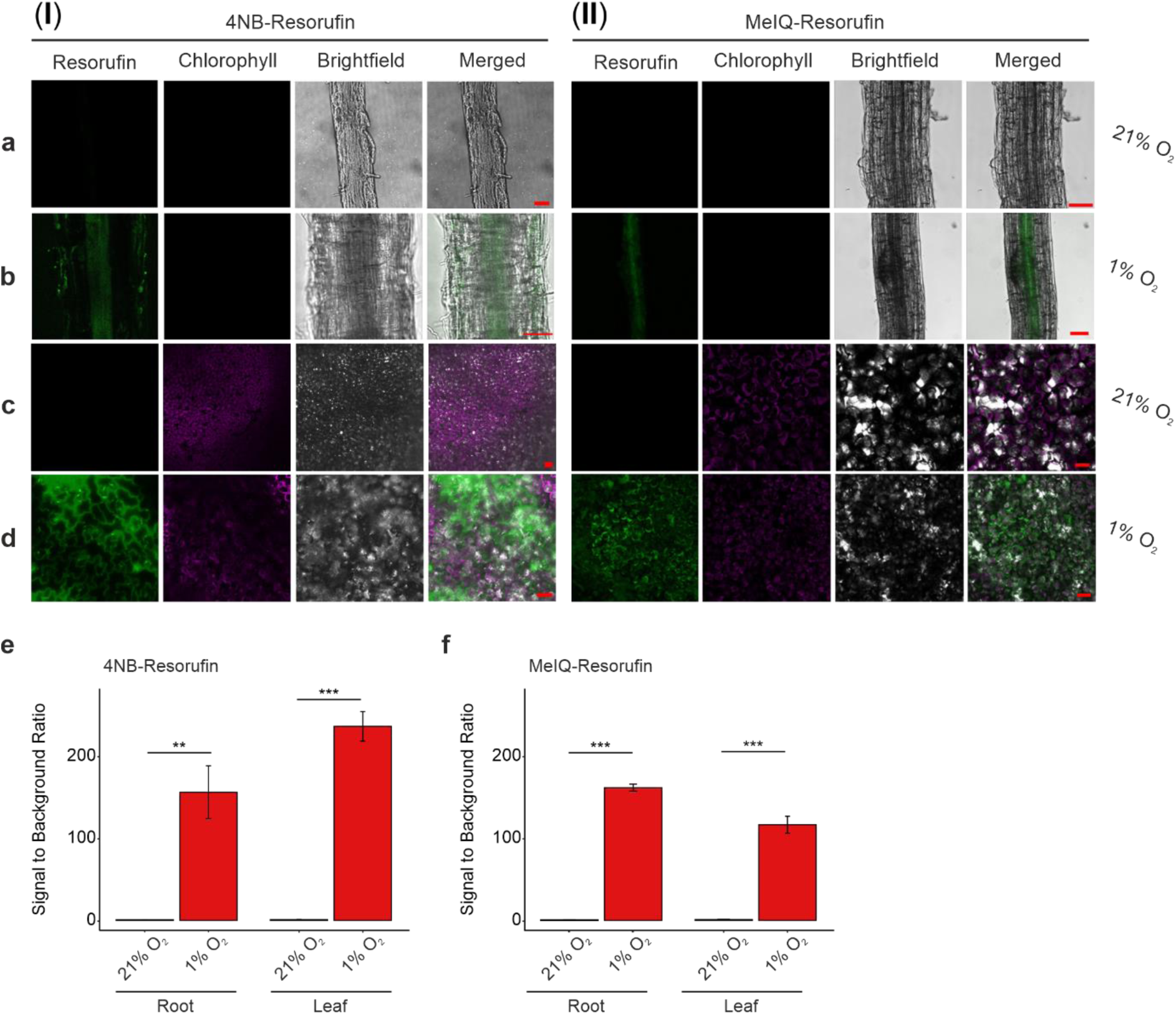
Bioreductive hypoxia responsive chemical probes show fluorescence in hypoxic tissue of Arabidopsis. Two-week old Arabidopsis seedlings were incubated with (**I**) 4NB-Resorufin or (**II**) MeIQ-Resorufin for 6 hours in normoxic (21 % O_2_) or hypoxic (1% O_2_) conditions. The live root (**a, b**) and leaf (**c, d**) of the seedlings were imaged using a confocal (Zeiss LSM880/LSM780) microscope using 561 nm laser to excite and fluorescence of the resorufin measured at 570 nm to 600 nm (green) as well as chlorophyll autofluorescence measured at 650 nm to 750 nm (purple). The leaf tissue experiments in panel (**I**) were performed with 50 µM 4NB-Resorufin, otherwise 100 µM probes used. Scale bars: 50 µm. (**e, f**) Quantification of resorufin fluorescence intensity in seedlings treated with 4NB- or MeIQ-Resorufin upon hypoxia or normoxia represented by the ratio of the signal intensity to the background (*n* = 3). Statistical significance was determined using two-sided Student’s t-test (** p-value < 0.01, *** p-value < 0.001).

To confirm that seedling intracellular O2 concentrations were hypoxic under the experimental hypoxia settings, we employed an *A. thaliana* line containing a luciferase reporter cassette under the control of the hypoxia responsive promoter element (HRPE). This HRPE is known to promote ERFVII-mediated gene upregulation under hypoxic conditions (Gasch *et al*., 2016; Akter *et al*., 2024). When seedlings were exposed to normoxia or hypoxia under the experimental conditions used for probe incubation, we detected a significant difference in luciferase activity, commensurate with elevated luciferase expression in hypoxia (Fig. **S7**), confirming that the seedlings were experiencing a hypoxic environment.

We next checked whether incubation of seedlings with the probes could affect plant growth. Seven-day old Arabidopsis seedlings were treated with 100 µM 4NB-Resorufin, MeIQ- Resorufin or resorufin (Fig. **S8**). Treatment with 10% DMSO was used as negative control. Physiological measurements were taken after four days of the treatment, during which seedlings were kept in 16:8 photoperiod conditions, in air. Consistent with our results for cell culture, we did not observe phenotypical differences between seedlings treated with the probes compared to DMSO treatment (Fig **S8a**). Additionally, no statistical differences were measured in primary root length, growth rate and fresh weight compared to control conditions (Fig **S8b, c, d**). These results suggest that 4NB-resorufin, MeIQ-resofurin and their reduced form does not affect biomass or root growth in Arabidopsis, in the conditions used in this study. Collectively, these results demonstrate that the probes are capable of acting as fluorescent markers of hypoxia in live plant tissue as well as in cells.

### MeIQ-Resorufin can detect endogenous hypoxia in Arabidopsis thaliana root lateral primordia

Finally, we sought to determine whether these probes could be used to detect physiologically relevant hypoxia in an otherwise normoxic environment. Arabidopsis root lateral primordia represent a physiological hypoxic niche, arising due to rapid cell division and high metabolism and demonstrated by localised ERF-VII stabilisation and hypoxic-response gene upregulation (Shukla *et al*., 2019). We tested whether MeIQ-Resorufin could detect hypoxia in root lateral primordia by incubating seven-day old Arabidopsis seedlings with 100 µM probe for 2 hours in either normoxic or hypoxic conditions. We selected to use MeIQ-Resorufin for this experiment given the higher levels of fluorescence observed for this probe in the whole cell cultures. Under normoxic conditions, MeIQ-Resorufin derived fluorescence was not typically observed in primary root tissue, as expected (Fig. **4**), while fluorescence was consistently observed in root lateral primordia regions at all stages of development (Fig. **5**, Fig. **S9**). When these tissues were incubated in hypoxia, fluorescence was usually observed in all regions of the root, including the lateral primordia (Fig. **5**, Fig. **S9**), consistent with previous experiments.. These results confirm that the MeIQ-Resorufin probe can be used to visualise regions of hypoxia in otherwise normoxic Arabidopsis tissues.

**Figure 5.**
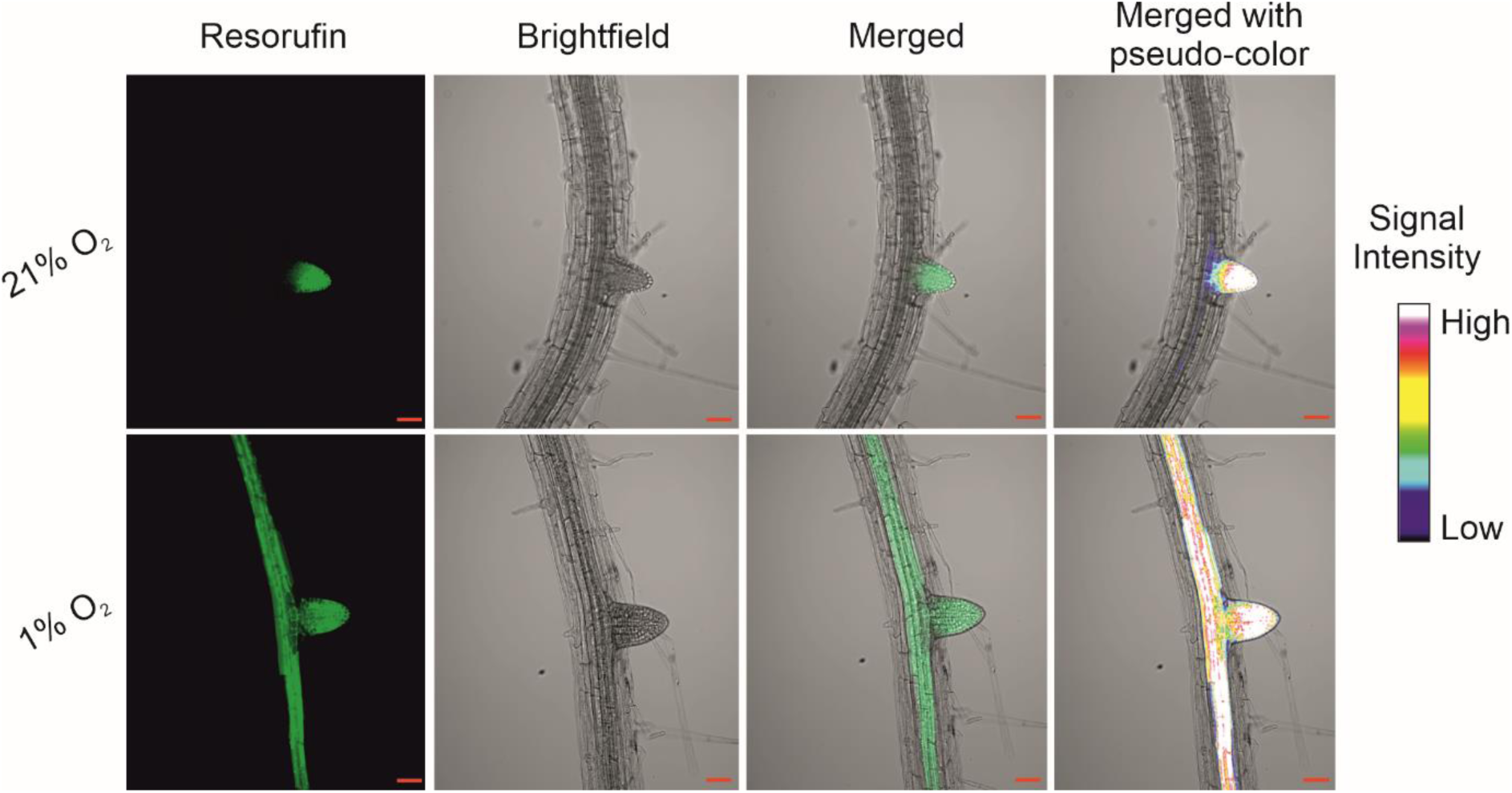
Bioreductive hypoxia responsive chemical probes show fluorescence in hypoxic tissue of Arabidopsis. Seven-day old seedlings were incubated with 100 µM MeIQ-Resorufin for 2 hours in normoxic (21 % O_2_ v/v) or hypoxic (1 % O_2_ v/v) environments. MeIQ-Resorufin fluorescence was imaged using a Zeiss LSM800 Airyscan laser scanning confocal microscope with 561 nm excitation laser light and 565-616 nm emission detection. Scale bars 50 µm; signal intensity bar refers to pseudocolour.

## Discussion

The hypoxia-sensing fluorescent probes 4NB-Resorufin and MeIQ-Resorufin are capable of being reduced under hypoxic conditions to release fluorescent resorufin, both *in vitro* and in bacterial and mammalian cells (Collins *et al*., 2018; Wallabregue *et al*., 2023). In this work, we have shown that these probes are also capable of being used to detect hypoxic conditions in plant cells, specifically the model plant *Arabidopsis thaliana*. Moreover, we were able to demonstrate that the probes can also be applied to *Arabidopsis thaliana* seedlings and fluoresce in exogenously applied hypoxic conditions (1 % O_2_) as well as endogenous hypoxia which exists in rapidly metabolising lateral root primordia. This work demonstrates that hypoxia-sensitive bioreductive enzymes must be present and active in the plant cells and tissues examined, similar to those in other kingdoms (Elmes, 2016; Cenas *et al*., 2021) in order to result in release of fluorescent resorufin.

Probe-derived fluorescence was visible through the use of confocal microscopy but also through the measurement of fluorescence intensity in a plate-based format. This format enables higher-throughput analysis of absolute fluorescence levels in a semi-quantitative manner, such that intensity is dependent on O_2_ availability. However, this was dependent on which probe was used: MeIQ- Resorufin derived fluorescence was significant at 5 % O_2_, in contrast to fluorescence derived from 4NB- Resorufin at the same O_2_ concentration. This indicates that MeIQ-Resorufin is able to detect ‘mild’ hypoxia (5 % O_2_ to 0.5 % O_2_) in contrast to 4NB-Resorufin which gave significant fluorescence values at 1 % O_2_ and below in planta. The mild hypoxia-dependent response of MeIQ resorufin is consistent with the hypoxia-sensitivity results observed for this probe in mammalian A549 lung cancer cells and HCT116 colorectal cancer cells spheroids (Wallabregue *et al*., 2023). In these systems MeIQ-Resorufin begins producing significant fluorescence at 4 % O₂, where a 5-fold increase compared to normoxic controls was observed after 2 hours incubation. Fluorescence increase further intensifies at lower oxygen concentrations with a maximum of 30-fold increase at <0.1 % O₂ compared to normoxic controls. In this study, the overall fluorescence levels derived from MeIQ-resorufin were approximately 5-fold higher than those from 4NB-resorufin. Time-course experiments conducted in *Arabidopsis thaliana* whole plants and cell lysates at 0.1 % O₂ further highlight these differences (Fig. **S5** and **S10**). Me-IQ-Resorufin exhibited a significant hypoxia-induced fluorescence increase, with 134-fold and 157- fold enhancements compared to the DMSO control in plant cell lysates with and without protease inhibitor cocktails, respectively, after 100 minutes. Similarly, a 196-fold increase was observed in whole plants after 360 minutes. In contrast, 4-NB-Resorufin showed only modest fluorescence increases under the same conditions, with 1.6-fold and 1.7-fold increases in cell lysates with and without protease inhibitor cocktails, respectively, and a 2-fold increase in whole plants compared to the DMSO control. Notably, the fluorescence increases between the two probes differed significantly, with 117- fold and 59-fold higher activation for Me-IQ-Resorufin in plant cell lysates without protease inhibitor cocktail and whole plants, respectively. These findings clearly demonstrate that the differences in activation are predominantly due to the inherently higher bioreductive reactivity of the indolequinone group compared to the 4-NB benzyl group, though a minor contribution from probe uptake cannot be excluded.

In hypoxic mammalian cells, bioreduction of MeIQ-resorufin is rapid (Wallabregue *et al*., 2023) which aligns with the observed higher fluorescence derived from MeIQ-resorufin than 4NB-Resorufin in hypoxic plant cells. Notably in cell cultures, the high levels of fluorescence observed with MeIQ- resorufin were not stable over time, with decreases in fluorescence being observed at low O_2_ (< 0.5 % O_2_) concentrations. This may be due to acid-quenching of resorufin fluorescence (Fig. **S4**), either in mildly acidic conditions associated with hypoxia in cytosol (Felle, 2005), or due to vacuolar xenobiotic compartmentalisation (Coleman *et al*., 1997), where pH levels will likely lead to a significant decrease in fluorescence. Despite the observation of this effect in whole cells, MeIQ-derived resorufin fluorescence showed prolonged stability in tissues which may reflect improved tolerance of hypoxia-induced metabolic acidosis in tissues compared to isolated whole cells (Felle, 2005). Importantly, the ability of MeIQ-resorufin to detect mild hypoxia with relatively high signal intensity enabled its use for the detection of endogenous hypoxia in lateral root primordia and may make this the most practical of the two probes for future use *in planta*.

The probes themselves are not significantly bioreduced in the presence of reactive oxygen species (Fig. **S11** and **S12**), which are known to be elevated in prolonged hypoxia (Jethva *et al*., 2023). However, they are susceptible to prolonged photobleaching and high concentrations of NaCl (Fig. **S13** and **S14**), so care should be taken in choosing conditions for their application. Nevertheless, the bioreductive ability of these chemical probes to detect hypoxic conditions in plant cells and tissue will be a valuable and timely addition to the suite of techniques available to measure O_2_ availability in the field of plant hypoxia research, both with respect to acute hypoxia (as an environmental stress) and chronic hypoxia (as a developmental cue). They can be applied without the requirement for genetic engineering of biosensors or the challenging aspects of probe measurement (including tissue damage and handling challenges) (Weits, 2021); determination of probe permeability and hypoxia-activation in crop plants will expand their utility. Furthermore, the inference that bioreductive enzymes are present and active in Arabidopsis cells indicates that the same principle or bioreduction could be used to trigger hypoxic activation of other molecules, a principle which has been exploited in mammalian hypoxia biology (Collins *et al*., 2018) and which may enable intervention in a range of situations where hypoxia is relevant.

## Resource availability

Data acquired in this study are available in the Supplementary Information or on request from the corresponding author. Methodology for probe synthesis is included in the Supplementary Information. Requests for resources (including 4NB-Resorufin and MeIQ-Resorufin) can be made to the corresponding author.

## Acknowledgements

The authors are grateful for funding from the UK Engineering and Physical Sciences Research Council (EPSRC) for the award of a Programme Grant to EMH and SJC (grant reference EP/S019901/1) that supported ALDW, HB, and EF. We thank the European Research Council (ERC) under the European Union’s Horizon 2020 research and innovation program (PCOMOD project, grant agreement 864888 (EF). We thank Prof. Nick Kruger (University of Oxford) for kind donation of the *A.thaliana* (c.v. Landsberg *erecta*) whole cell culture.

## Author contributions

MP, MSK, ALDW, VV and AMR conducted experiments and analysed data, assisted by ENS and HB. EF, SJC and EMH conceived the research. EF and DW designed and supervised the research. SJC supervised the chemistry aspects of the research. MP, MSK, ALDW, and EF wrote the manuscript with assistance from all co-authors. MP, MSK and ALDW contributed equally to this work.

## Completing interest

The authors declare that they have no conflicts of interest with the contents of this article.

