## Supplementary Information for "Hypoxia-activated fluorescent probes as markers of oxygen levels in plant cells and tissues"

Monica Perri[1,†], M. Shahneawz Khan[1,2,†], Antoine L. D. Wallabregue[2†], Viktoriia Voloboeva[3,4], Amber M. Ridgway[2,5], Edward N. Smith[1], Hannah Bolland[6,7], Ester M. Hammond[6], Stuart J. Conway[2,8], Daan A. Weits[3], Emily Flashman[*2]

[1] Department of Biology, University of Oxford, South Parks Road, Oxford, OX1 3RB, U.K.

[2] Department of Chemistry, Chemistry Research Laboratory, University of Oxford, Mansfield Road, Oxford, OX1 3TA, U.K.

[3] Experimental and Computational Plant Development, Institute of Environment Biology, Utrecht University, Padualaan 8, Utrecht 3584 CH, Netherlands

[4] PlantLab, Institute of Plant Sciences, Scuola Superiore Sant'Anna, 56010 Pisa, Italy

[5] Current address: Department of Biology and Epigenetics Institute, University of Pennsylvania, Philadelphia, PA 19104-4544, USA.

[6] Department of Oncology, University of Oxford, Old Road Campus Research Building, Oxford, OX3 7DQ, U.K.

[7] Current address: School of Veterinary Medicine, Faculty of Health and Medical Sciences, University of Surrey, Guildford, Surrey, GU2 7AL, U.K.

[8] Current address: Department of Chemistry & Biochemistry, University of California Los Angeles, 607 Charles E. Young Drive East, P. O. Box 951569, Los Angeles, California, 90095-1569, USA.

[†] Authors contributed equally

The following Supporting Information is available for this article:

**Figure S1.** Additional images of *A. thaliana cells* incubated with 4NB-Resorufin and MeIQ-Resorufin.

**Figure S2.** Fluorescence at 570-590 nm is resorufin-dependent and conditionals on cellular uptake.

**Figure S3.** 4NB-Resorufin and MeIQ-Resorufin are not toxic to *Arabidopsis thaliana* cells and seedlings under the conditions used.

**Figure S4.** Resorufin, MeIQ-Resorufin, and 4NB-Resorufin fluorescence is quenched at mildly acidic pH.

**Figure S5.** MeIQ-Resorufin and 4NB-Resorufin bioreduction in *A. thaliana* seedling under hypoxia.

**Figure S6.** *A. thaliana* seedlings elicit a typical hypoxic response after 1 hour incubation in hypoxia (1% $O_2$).

**Figure S7.** 4NB-Resorufin, MeIQ-Resorufin and their reduced form Resorufin do not affect growth in Arabidopsis seedlings.

**Figure S8.** Hypoxic fluorescence in *A. thaliana* tissues is a result of incubation with 4NB-/MeIQ-Resorufin.

**Figure S9.** Additional images of MeIQ-Resorufin-derived fluorescence in *A. thaliana* lateral root primordia, including at different stages of development .

**Figure S10**. MeIQ-Resorufin and 4NB-Resorufin bioreduction in *A. thaliana* cell lysates under hypoxia.

**Figure S11**. Stability of 4NB-Resorufin under treatment with hydrogen peroxide or hydroxyl radical.

**Figure S12**. Stability of MeIQ-Resorufin under treatment with hydrogen peroxide or hydroxyl radical.

**Figure S13**. Resorufin, MeIQ-Resorufin, and 4NB-Resorufin fluorescence is quenched by after light irradiation.

**Figure S14**. Resorufin, MeIQ-Resorufin, and 4NB-Resorufin fluorescence is quenched in buffer with high salt concentrations.

**Chemistry Experimental Details** General experimental techniques plus different *in vitro* assays, synthetic procedures and HPLC chromatograms for preparation of 4NB-Resorufin and MeIQ-Resorufin.

**References**

**Figure S1. Additional images of *A. thaliana* cells incubated with 4NB-resorufin and MeIQ-resorufin**. *A. thaliana* cell cultures were incubated with 10 µM of either 4NB-resorufin or MeIQ-resorufin under normoxic (21 % $O_2$) or hypoxic (1 % $O_2$) conditions for 2 hours. Fluorescence signals were collected for resorufin (green, 570-590 nm) and chlorophyll autofluorescence (purple, 650-750 nm). Scale bars: 20 µm.

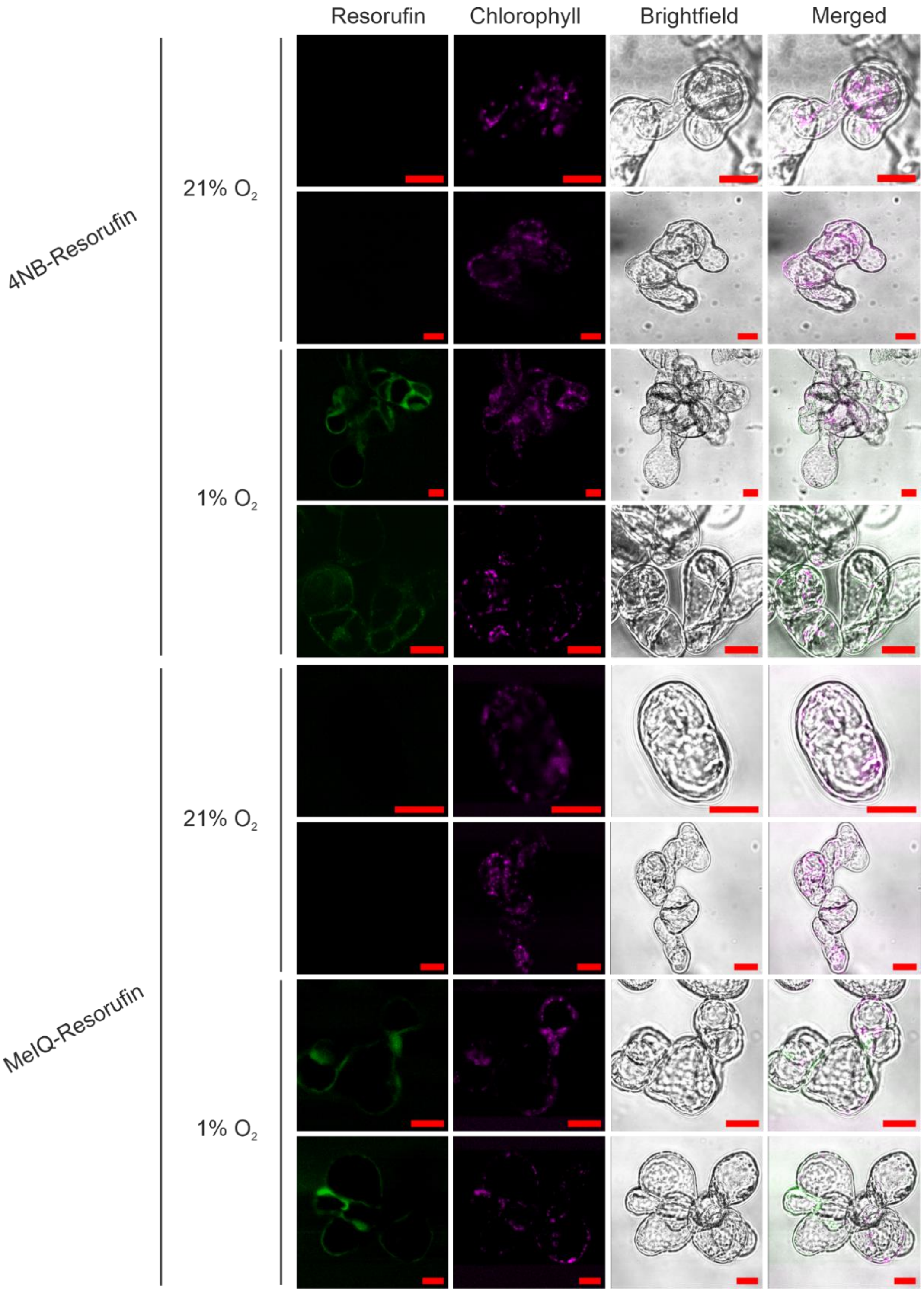

**Figure S2. Fluorescence at 570-590 nm is resorufin-dependent and conditional on cellular uptake.** Confocal images demonstrating that fluorescence observed in hypoxic *A. thaliana* cells is dependent on the presence of 4NB or MeIQ-Resorufin and its ability to penetrate cells. Four-day old *A. thaliana* whole cell cultures were incubated for 2 hours with 1 % DMSO (**a**) under normoxic (21 % $O_2$) or hypoxic conditions (1 % $O_2$). MS media (in the absence of cells) was incubated with 10 µM of either 4NB-resorufin (**b**) or MeIQ-resorufin (**c**) in normoxia or hypoxia, respectively. Scale bars: 20 µm.

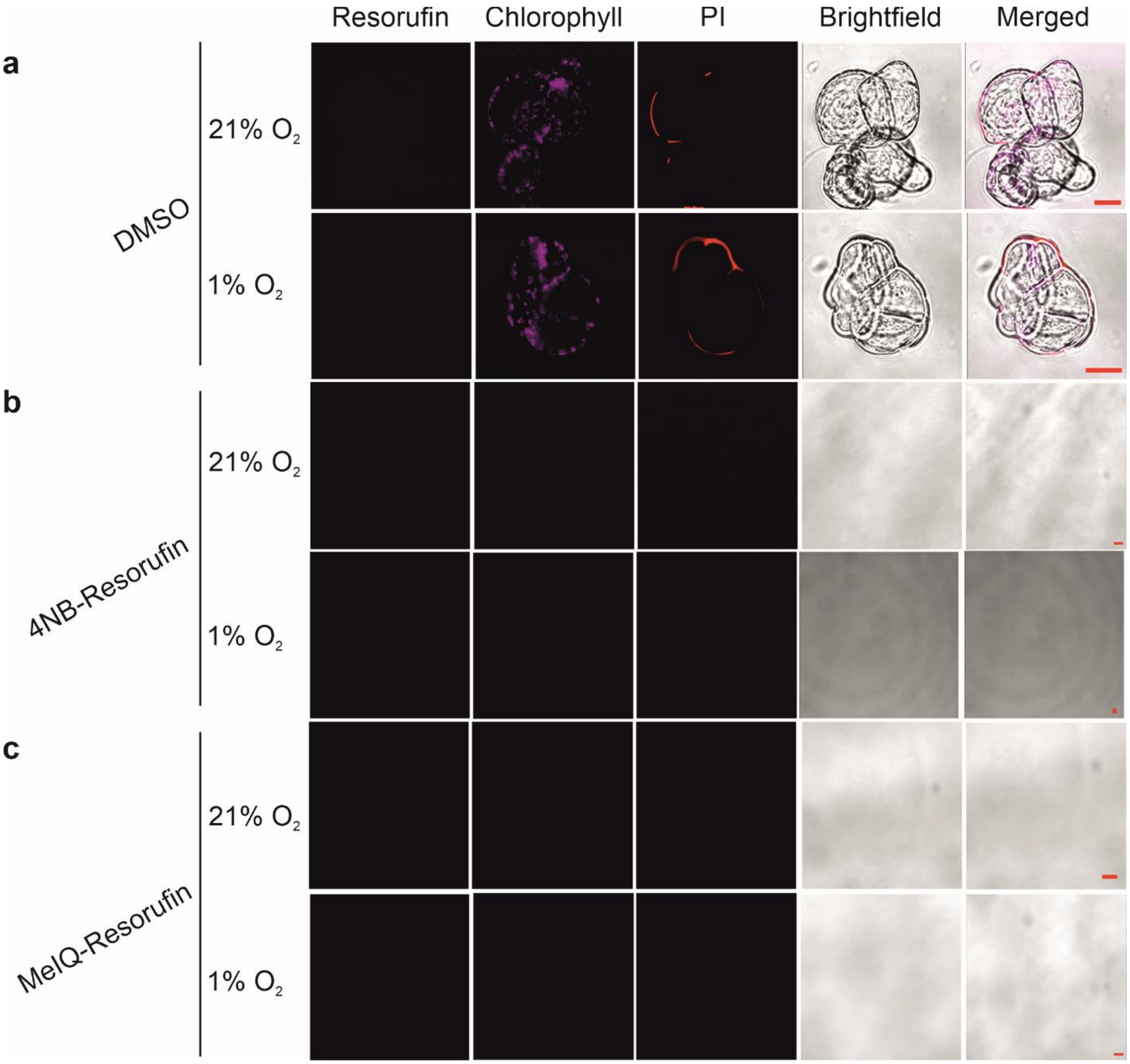

**Figure S3. 4NB-resorufin and MeIQ-resorufin are not toxic to *Arabidopsis thaliana* cells under the conditions used.** Arabidopsis cells (four-day old) in MS media (1 mL) were treated with varying concentrations (10, 50 and 100 µM) of 4NB-Resorufin (**a**) or MeIQ-Resorufin (**b**) in normoxia (21 % $O_2$) or hypoxia (1 % $O_2$) for 2 hours followed by propidium iodide (PI) staining by confocal imaging of the cells. DMSO was used as a control for each concentration. Cells stained internally with PI dye were considered dead cells. Dead cells were scored from triplicate slide images within a fixed area (0.83 µm x 0.83 µm) taken at 20x magnification using Zeiss LSM780. Letters indicate statistical differences between conditions analysed using two-way ANOVA, followed by Tukey HSD *post-hoc* test (p-value < 0.05).

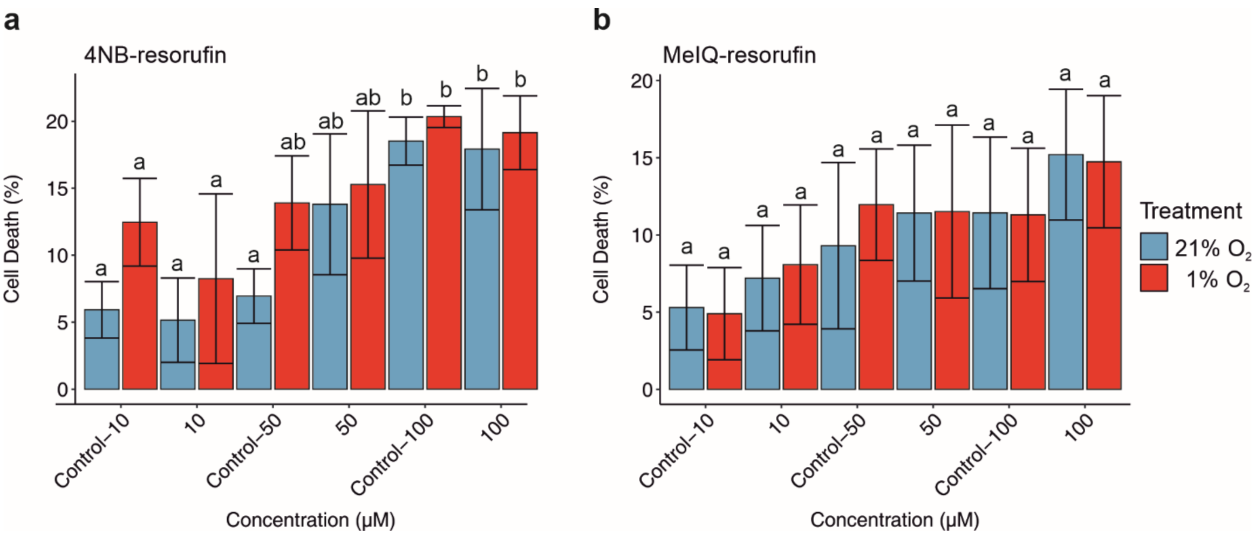

**Figure S4. Resorufin fluorescence is quenched at mildly acidic pH.** (**a**) 3000 µL of potassium phosphate buffer (500 mM) with adjusted pH (5 or 7.4) was mixed with resorufin (10 µM) in a quartz cuvette. The fluorescence emission was measured from 560 to 700 nm (excitation 545 nm, slits 3, 3 nm) under ambient conditions. (**b**) 200 µL of sodium phosphate buffer (200 mM) with adjusted pH (4 - 8.9) was mixed with resorufin (10 µM). The plate was agitated orbitally for 30 s at 100 rpm for homogenous mixing before measuring the fluorescence emission at 580 nm (excitation, 550 nm). Data represents mean of triplicate measurements ($n$ = 3) and the vertical bar represents standard deviations of the measurements. Letters indicate statistical differences between conditions analysed using one-way ANOVA, followed by Tukey HSD *post-hoc* test (p-value < 0.05). (**c**) Resorufin (1 µM) in Milli-Q water adjusted to the indicated pH, following the general procedure. (**d**) Quantification of the average fluorescence intensity from three replicates shows a decrease in resorufin fluorescence at pH 5, where the fluorescence intensity at pH 7.4 is normalized to 100 %. Statistical significance was assessed using one-way ANOVA, followed by Tukey HSD *post-hoc* test (p-value < 0.05), where letters indicate statistical differences between groups. (**e**) MeIQ-Resorufin (1 µM) in Milli-Q water adjusted to the indicated pH, following the general procedure. (**f**) Quantification of the average fluorescence intensity from three replicates shows a decrease in MeIQ-Resorufin fluorescence at pH 5, with the intensity at pH 7.4 normalized to 100%. Statistical significance was assessed using two-sided Student's t-test (***p-value < 0.001). (**g**) 4NB-Resorufin (1 µM) in Milli-Q water adjusted to the indicated pH, following the general procedure. Fluorescence intensity data represent the mean of three replicates, collected with excitation at 545 nm and slit widths set to 3 nm for both excitation and emission. (**h**) Quantification of the mean fluorescence intensity from three replicates shows a decrease in 4NB-Resorufin fluorescence at pH 5, where the fluorescence intensity at pH 7.4 is normalized to 100%. Statistical significance was assessed using two-sided Student's t-test (***p-value < 0.001).

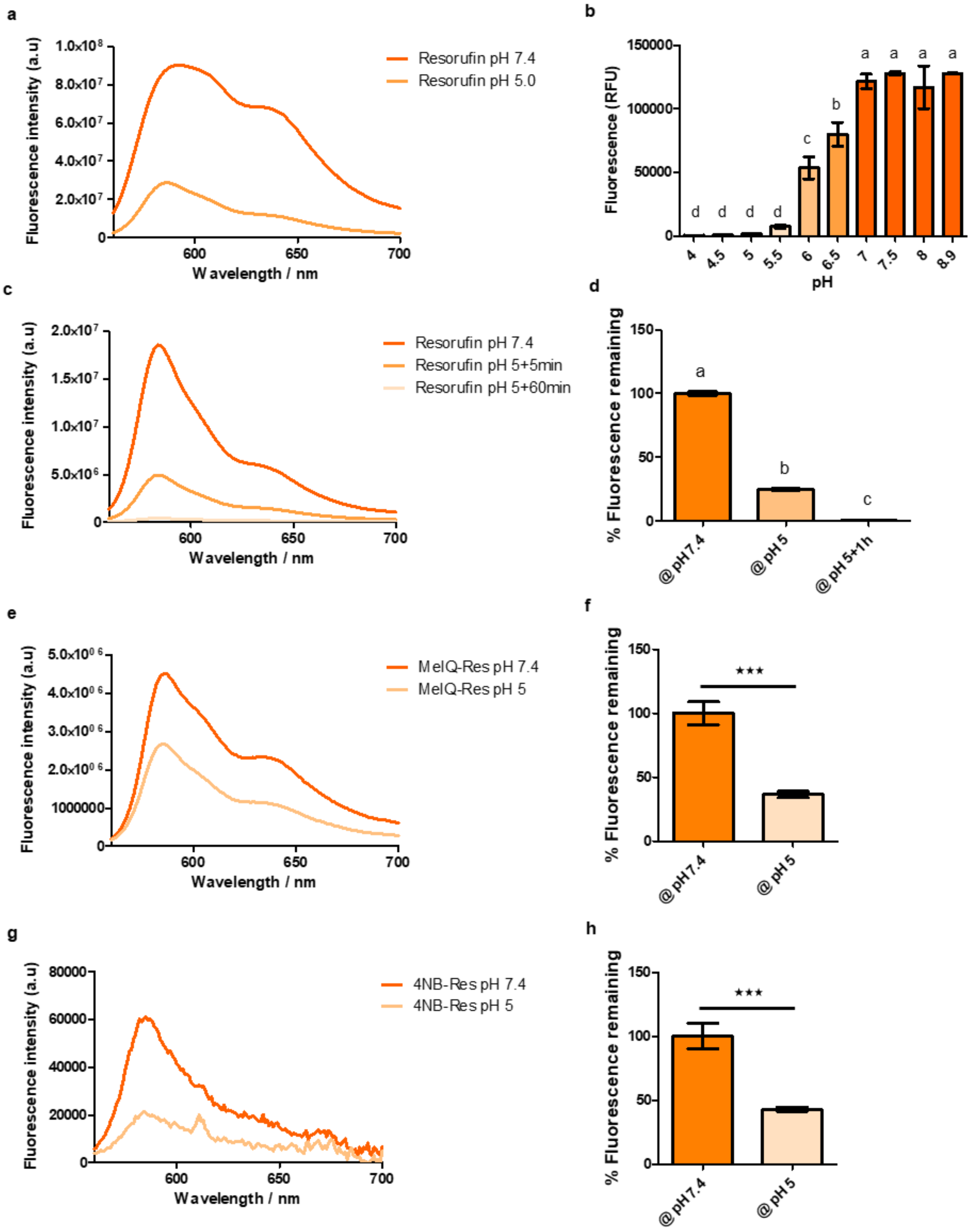
a
Fluorescence intensity (a.u)
1.0x10⁸
8.0x10⁷
6.0x10⁷
4.0x10⁷
2.0x10⁷
0
600
650
700
Wavelength / nm
Resorufin pH 7.4
Resorufin pH 5.0
b
Fluorescence (RFU)
150000
100000
50000
0
d d d d c b a a a a
4 4.5 5 5.5 6 6.5 7 7.5 8 8.9
pH
c
Fluorescence intensity (a.u)
2.0x10⁷
1.5x10⁷
1.0x10⁷
5.0x10⁶
0
600
650
700
Wavelength / nm
Resorufin pH 7.4
Resorufin pH 5+5min
Resorufin pH 5+60min
d
% Fluorescence remaining
150
100
50
0
a
b
c
@pH7.4
@pH5
@pH5+1h
e
Fluorescence intensity (a.u)
5.0x10⁰⁶
4.0x10⁰⁶
3.0x10⁰⁶
2.0x10⁰⁶
1000000
0
600
650
700
Wavelength / nm
MeIQ-Res pH 7.4
MeIQ-Res pH 5
f
% Fluorescence remaining
150
100
50
0
***
@pH7.4
@pH5
g
Fluorescence intensity (a.u)
80000
60000
40000
20000
0
600
650
700
Wavelength / nm
4NB-Res pH 7.4
4NB-Res pH 5
h
% Fluorescence remaining
150
100
50
0
***
@pH7.4
@pH5

**Figure S5. MeIQ-Resorufin and 4NB-Resorufin bioreduction in *Arabidopsis thaliana* seedlings at 0.1 % $O_2$.** (**a**, **b**) Proposed reactions of MeIQ-Resorufin and 4NB-Resorufin with Arabidopsis seedlings leading to the formation of resorufin. (**c**, **d**) Hypoxic incubation of whole Arabidopsis seedlings (10-day old) in 500 µL of Milli-Q water containing 100 µM of MeIQ-Resorufin (**c**) or 4NB-Resorufin (**d**) was performed for 360 minutes at 25 °C. The mean fluorescence intensity of three biological replicates measured after 360 minutes, DMSO was used as a control for each concentration. (**e**) Quantification of the fluorescence increase for MeIQ-Resorufin observed in panel c (where the fluorescence is normalized to the DMSO control). (**f**) Quantification of the fluorescence increase for 4NB-Resorufin observed in panel d (where the fluorescence is normalized to the DMSO control). (**g**) Mean fluorescence intensity of three biological replicates measured after 360 minutes reveals a difference in the rate at which maximum intensity is achieved for MeIQ-Resorufin (dashed line) compared to 4NB-Resorufin (solid line). (**h**) Quantification of the fluorescence increase for MeIQ-Resorufin and 4NB-Resorufin observed in panel g. Data are representative of three biological replicates; statistical significance was assessed using two-sided Student’s t-test (***p-value < 0.001).

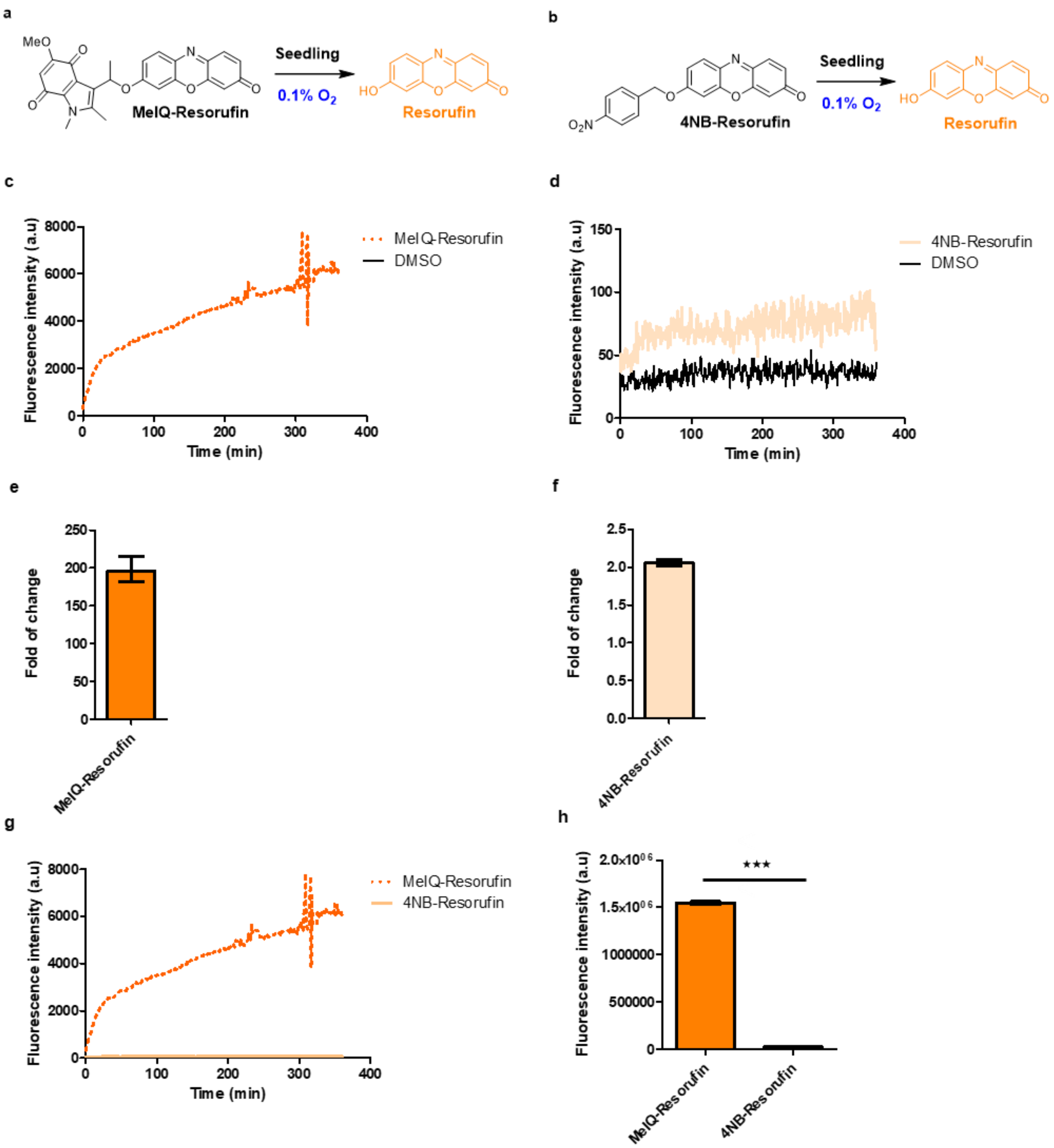
a
MeO
MeIQ-Resorufin
Seedling
0.1% O2
HO
Resorufin
b
O2N
4NB-Resorufin
Seedling
0.1% O2
HO
Resorufin
c
Fluorescence intensity (a.u)
Time (min)
MeIQ-Resorufin
DMSO
d
Fluorescence intensity (a.u)
Time (min)
4NB-Resorufin
DMSO
e
Fold of change
MeIQ-Resorufin
f
Fold of change
4NB-Resorufin
g
Fluorescence intensity (a.u)
Time (min)
MeIQ-Resorufin
4NB-Resorufin
h
Fluorescence intensity (a.u)
***
MeIQ-Resorufin
4NB-Resorufin

**Figure S6. Hypoxic fluorescence in *Arabidopsis thaliana* tissues is a result of incubation with 4NB-/MeIQ-Resorufin.** *A. thaliana* seedlings were floated in media containing an equivalent volume of DMSO to that used in treatment with 4NB-/MeIQ-Resorufin (50 µL/100 µL, respectively). No fluorescence was observed under either normoxia or hypoxia in roots (**a, b**) or leaf tissue (**c, d**). Scale bars: 50 µm.

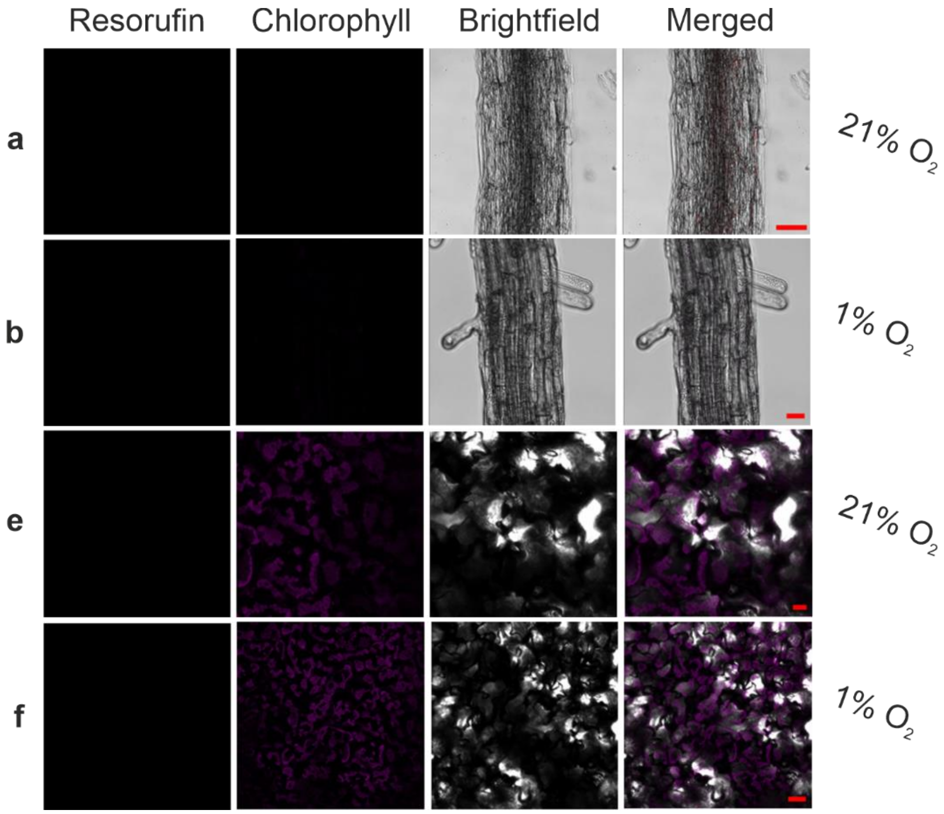

**Figure S7.** ***Arabidopsis thaliana*** **seedlings elicit a typical hypoxic response after 1 hour incubation in hypoxia (1% $O_2$).** *A. thaliana* wild-type seedlings expressing a hypoxia-responsive cassette comprising hypoxia-response element (HRPE) and UTR-ADH promoter upstream of a gene encoding firefly luciferase enzyme. When a plant experiences hypoxia, hypoxia-responsive transcription factors recognize the HRPE motif and upregulate downstream genes (Gasch *et al.*, 2016). Luciferase expression therefore indicates the seedlings are experiencing hypoxia. Two-weeks old seedlings were incubated in 1 mL MS media for 1 hour in the Clariostar plate reader coupled to the atmospheric control unit, either in hypoxia (1 % $O_2$) or normoxia (21 % $O_2$). A vivazine™-based live assay system was employed to determine luminescence arising from luciferase activity. Data are representative of four biological replicates, statistical significance was assessed using Kruskal-Wallis, followed by Dunn's test. Seedlings exposed to hypoxia showed significantly higher levels of luciferase activity (*** p-value < 0.001).

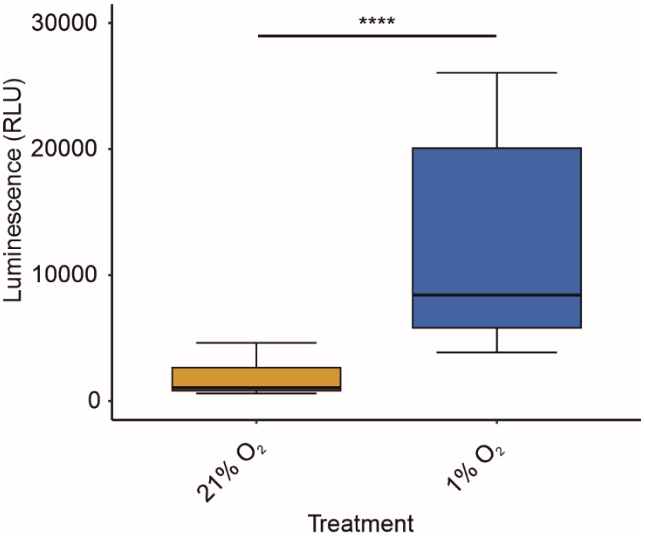

**Figure S8. 4NB-Resorufin, MeIQ-Resorufin and their reduced form resorufin do not affect growth in Arabidopsis seedlings.** (**a**) Phenotypes of seven-day old Arabidopsis seedlings, after 4 days of treatment with 10 % DMSO, 100 µM 4NB-resorufin, MeIQ-resorufin or resorufin. Scale bar: 1 cm. (**b**, **c**, **d**) Measurements of growth rate per day, primary root length and fresh weight at the end of the treatment ($n \geq 21$). Letters indicate statistical differences between conditions analysed using one-way ANOVA, followed by Tukey HSD *post-hoc* test.

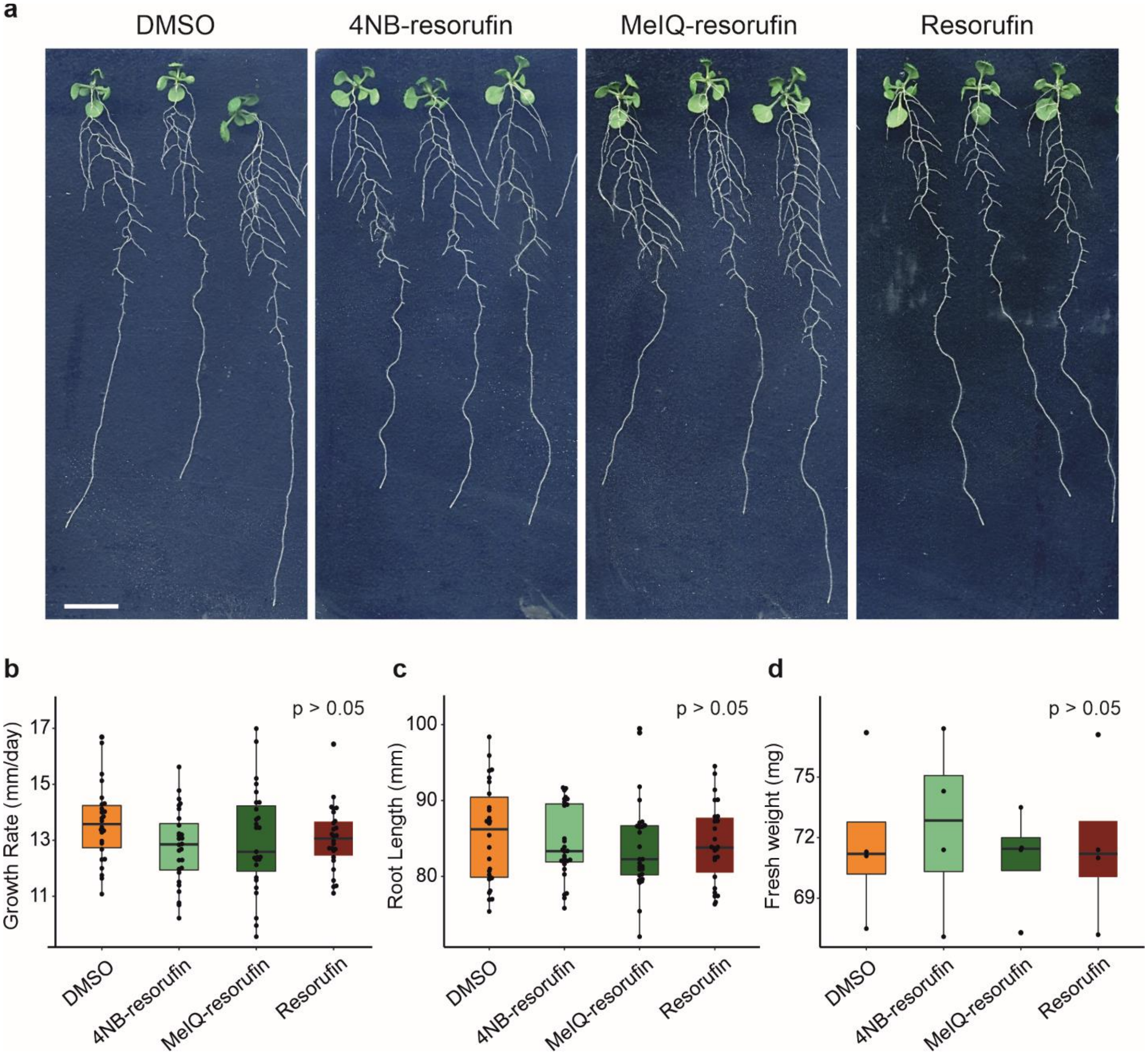

**Figure S9. Additional images of MeIQ-Resorufin-derived fluorescence in *Arabidopsis thaliana* lateral root primordia, including at different stages of development.** In three separate experiments at 21 % $O_2$, 14 lateral roots from a total of four plants were imaged (10 shown below). The hypoxic environment in the lateral root primordia (Shukla *et al.*, 2019) is shown by intensity of fluorescence (indicated by pseudo-colour); lateral root primordia at a range of stages of development were shown to be hypoxic. In 2/14 cases (for example #1) a fluorescent signal was also seen in the root, however in these cases signal intensity was less in the main root than in the lateral root primordia. In two separate experiments at 1 % $O_2$, 16 lateral roots from a total of four plants were imaged (10 shown below). In most cases, fluorescent signal was observed in both lateral roots and the main root, however in 2/16 cases (for example #2), fluorescence was only observed in the lateral root. Scale bars = 50 µm.

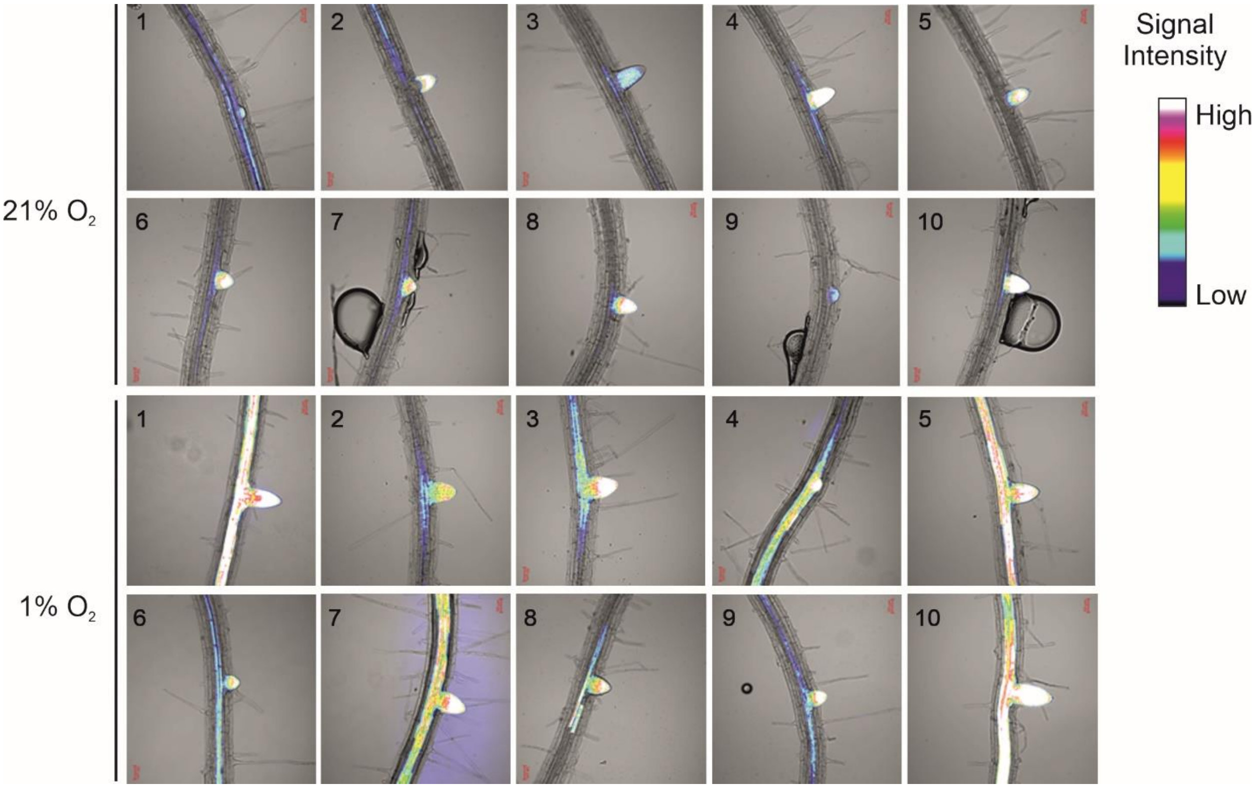

**Figure S10. MeIQ-Resorufin gets bioreduced faster than 4NB-Resorufin at 0.1 % $O_2$ concentration in Arabidopsis thaliana lysates.** (**a**, **b**) Proposed reactions of MeIQ-Resorufin and 4NB-Resorufin with Arabidopsis lysates leading to the formation of resorufin. (**c**, **d**) Hypoxic incubation of Arabidopsis lysates (10 days old, 147 µL) with or without protease inhibitor cocktail (PIC), NADH solution (1.5 µL of a 10 mM stock solution), and 10 µM of either MeIQ-Resorufin (**c**) or 4NB-Resorufin (**d**) was conducted for 100 minutes at 25 °C. Average fluorescence intensity of three biological replicates measured after 360 minutes, DMSO was used as a control for each concentration. (**e**) Quantification of the fluorescence increase for MeIQ-Resorufin observed in panel c (where the fluorescence is normalized to the DMSO control). (**f**) Quantification of the fluorescence increase for 4NB-Resorufin observed in panel d (where the fluorescence is normalized to the DMSO control). (**g**) Average fluorescence intensity of three biological replicates measured after 100 minutes reveals a difference in the rate at which maximum intensity is achieved for MeIQ-Resorufin (dashed line) compared to 4NB-Resorufin (solid line). Statistical significance was assessed using two-sided Student's t-test (***p-value < 0.001). (**h**) Quantification of the fluorescence increase for MeIQ-Resorufin and 4NB-Resorufin observed in panel g. Data are representative of three biological replicates; statistical significance was assessed using one-way ANOVA, followed by Tukey HSD *post-hoc* test, where letters indicate statistical differences between groups (p-value < 0.05).

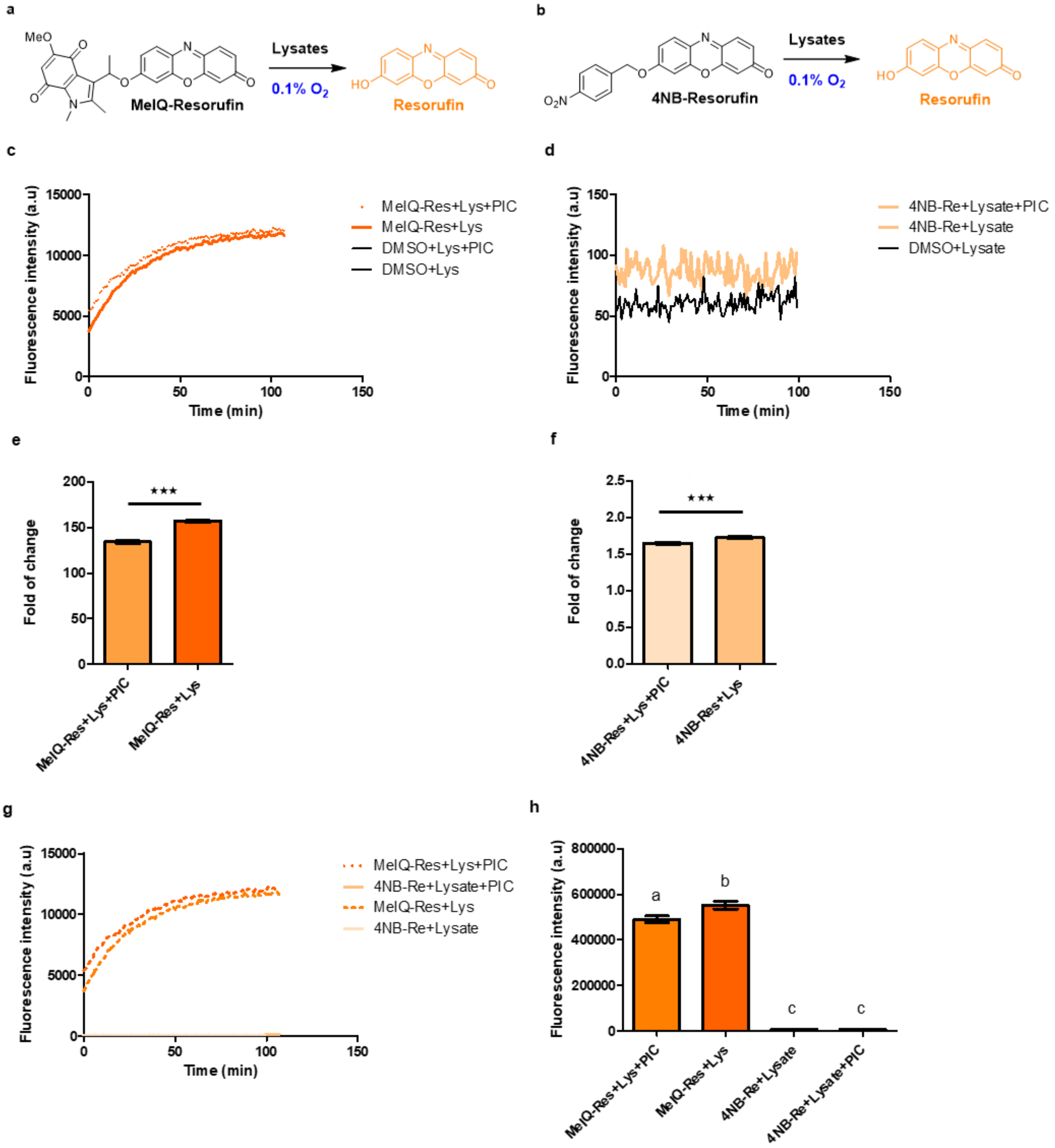
a
MeO
MeIQ-Resorufin
Lysates
0.1% O2
HO
Resorufin
b
O2N
4NB-Resorufin
Lysates
0.1% O2
HO
Resorufin
c
Fluorescence intensity (a.u)
Time (min)
MeIQ-Res+Lys+PIC
MeIQ-Res+Lys
DMSO+Lys+PIC
DMSO+Lys
d
Fluorescence intensity (a.u)
Time (min)
4NB-Re+Lysate+PIC
4NB-Re+Lysate
DMSO+Lysate
e
Fold of change
***
MeIQ-Res+Lys+PIC
MeIQ-Res+Lys
f
Fold of change
***
4NB-Res+Lys+PIC
4NB-Res+Lys
g
Fluorescence intensity (a.u)
Time (min)
MeIQ-Res+Lys+PIC
4NB-Re+Lysate+PIC
MeIQ-Res+Lys
4NB-Re+Lysate
h
Fluorescence intensity (a.u)
a
b
c
c
MeIQ-Res+Lys+PIC
MeIQ-Res+Lys
4NB-Re+Lysate
4NB-Re+Lysate+PIC

**Figure S11. Treatment of 4NB-Resorufin with hydrogen peroxide or hydroxyl radical does not result in the release of resorufin.** (**a**) The proposed reaction mechanism of 4NB-Resorufin with hydrogen peroxide or hydroxyl radicals predicts the formation of resorufin and corresponding nitrobenzyl derivatives. However, no evidence of resorufin or nitrobenzyl derivatives was observed. (**b**) 4NB-Resorufin (1 µM) in Milli-Q water (pH 7) was treated with hydrogen peroxide (100 eq.) or hydroxyl radicals (100 eq.) for 10 minutes, following the general procedure. Fluorescence intensity data are the mean fluorescence intensity of three biological replicates and were collected with excitation at 545 nm and slit widths set to 3 nm for both excitation and emission. (**c**) Quantification of the mean fluorescence intensity of three biological replicates shows no increase after 10 minutes of treatment with hydrogen peroxide or hydroxyl radical, indicating the stability of 4NB-Resorufin (where the fluorescence of untreated resorufin is normalized to 100 %). Data are representative of three biological replicates; statistical significance was assessed using one-way ANOVA, followed by Tukey HSD *post-hoc* test, where letters indicate statistical differences between groups (p-value < 0.05).

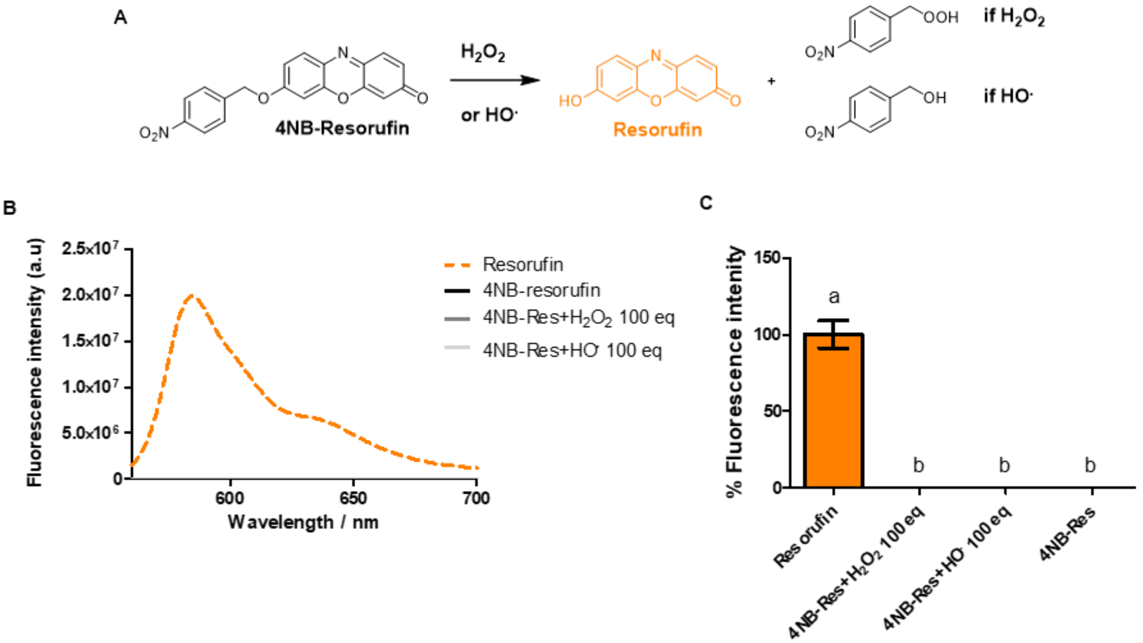

**Figure S12. Treatment of MeIQ-Resorufin with hydrogen peroxide or hydroxyl radical results in only a modest release of resorufin in the former case.** (**a**) The proposed reaction of MeIQ-Resorufin with hydrogen peroxide or hydroxyl radicals predicts the formation of resorufin and corresponding nitrobenzyl derivatives. (**b**) MeIQ-Resorufin (1 µM) in Milli-Q water (pH 7) was treated with hydrogen peroxide (100 eq.) or hydroxyl radicals (100 eq.) for 10 minutes, following the general procedure. Fluorescence intensity data are the mean fluorescence intensity of three biological replicates and were collected with excitation at 545 nm and slit widths set to 3 nm for both excitation and emission. (**c**) Quantification of the mean fluorescence intensity of three biological replicates shows a 24 % increase after 10 minutes of treatment with hydrogen peroxide, indicating modest release of resorufin (where the fluorescence of untreated resorufin is normalized to 100 %). Data are representative of three biological replicates; statistical significance was assessed using one-way ANOVA, followed by Tukey HSD *post-hoc* test, where letters indicate statistical differences between groups (p-value < 0.05).

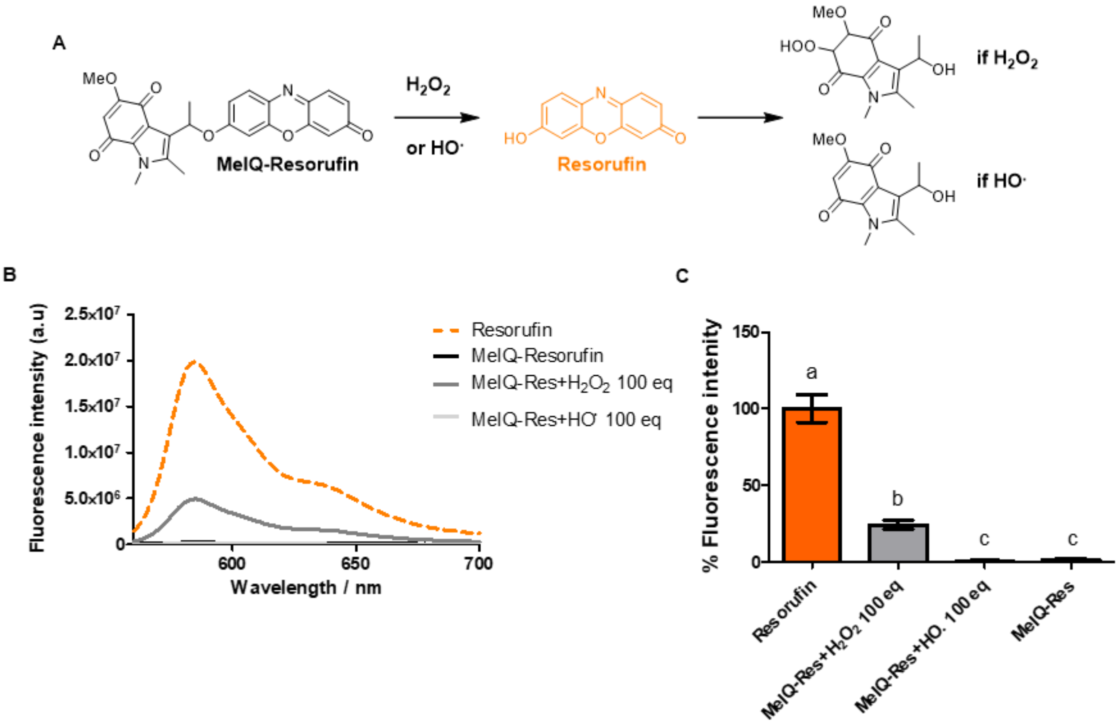

**Figure S13. Fluorescence of MeIQ-Resorufin, 4NB-Resorufin, and Resorufin is quenched after illumination with white LED (Photobleaching).** (**a**) MeIQ-Resorufin (1 µM) in Milli-Q water (pH 7) and exposed to light as described in the general procedure. (**b**) 4NB-Resorufin (1 µM) in Milli-Q water (pH 7) and exposed to light as described in the general procedure. (**c**) Resorufin (1 µM) in Milli-Q water (pH 7) and exposed to light as described in the general procedure. Fluorescence intensity data represent the mean of three replicates and were collected at the time points specified in the general procedure with excitation at 545 nm and slit widths set to 3 nm for both excitation and emission. (**d**) Quantification of the mean fluorescence intensity from three replicates shows a decrease in MeIQ-Resorufin fluorescence after 4 hours (T = 4 hours), normalized to the fluorescence of untreated MeIQ-Resorufin at the initial time point (T = 0, set as 100 %). (**e**) Quantification of the mean fluorescence intensity from three replicates shows a decrease in 4NB-Resorufin fluorescence after 4 hours (T = 4 hours), normalized to the fluorescence of untreated 4NB-Resorufin at the initial time point (T = 0, set as 100 %). (**f**) Quantification of the mean fluorescence intensity from three replicates shows a decrease in Resorufin fluorescence after 4 hours (T = 4h), normalized to the fluorescence of untreated Resorufin at the initial time point (T = 0, set as 100%). Statistical significance was assessed using two-sided Student's t-test (***p-value < 0.001).

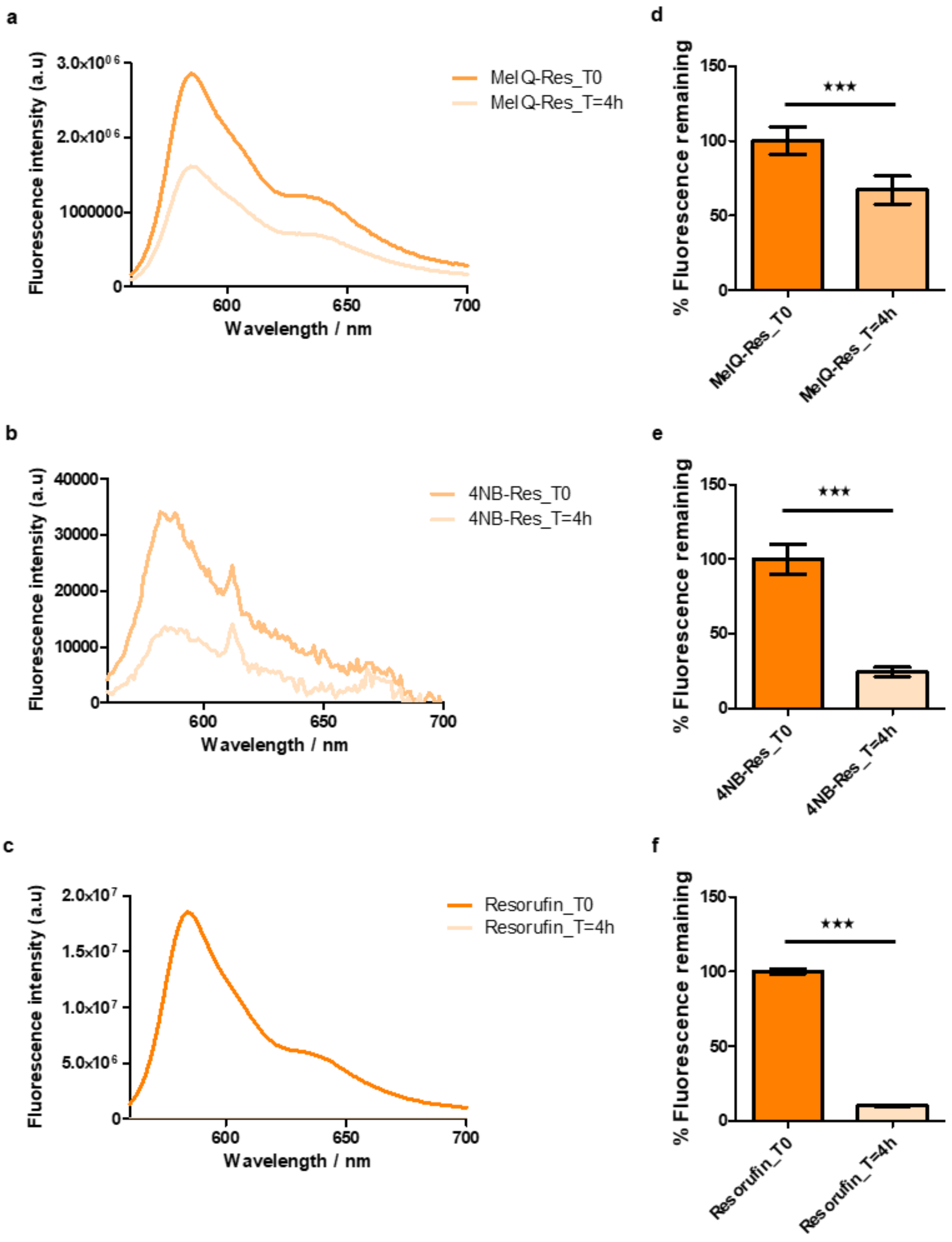
a
MeIQ-Res_T0
MeIQ-Res_T=4h
Fluorescence intensity (a.u)
3.0x10⁰⁶
2.0x10⁰⁶
1000000
0
600
650
700
Wavelength / nm
d
% Fluorescence remaining
150
100
50
0
***
MeIQ-Res_T0
MeIQ-Res_T=4h
b
4NB-Res_T0
4NB-Res_T=4h
Fluorescence intensity (a.u)
40000
30000
20000
10000
0
600
650
700
Wavelength / nm
e
% Fluorescence remaining
150
100
50
0
***
4NB-Res_T0
4NB-Res_T=4h
c
Resorufin_T0
Resorufin_T=4h
Fluorescence intensity (a.u)
2.0x10⁷
1.5x10⁷
1.0x10⁷
5.0x10⁶
0
600
650
700
Wavelength / nm
f
% Fluorescence remaining
150
100
50
0
***
Resorufin_T0
Resorufin_T=4h

**Figure S14.** Fluorescence of Resorufin, MeIQ-Resorufin, and 4NB-Resorufin is quenched in buffer with varying concentration of NaCl. (**a**) Resorufin (1 μM) in aqueous buffer containing different concentrations of NaCl, as described in the general procedure. (**b**) MeIQ-Resorufin (1 μM) in aqueous buffer containing different concentrations of NaCl, as described in the general procedure. (**c**) 4NB-Resorufin (1 μM) in aqueous buffer containing different concentrations of NaCl, as described in the general procedure. Fluorescence intensity data represent the average of three replicates, collected with excitation at 545 nm and slit widths set to 3 nm for both excitation and emission. (**d**) Quantification of the mean fluorescence intensity from three replicates shows a decrease in Resorufin fluorescence at different salt concentrations, normalized to the fluorescence of untreated Resorufin (set as 100 %). (**e**) Quantification of the mean fluorescence intensity from three replicates shows a decrease in MeIQ-Resorufin fluorescence at different salt concentrations, normalized to the fluorescence of untreated MeIQ-Resorufin (set as 100 %). (**f**) Quantification of the mean fluorescence intensity from three replicates shows a decrease in 4NB-Resorufin fluorescence at different salt concentrations, normalized to the fluorescence of untreated 4NB-Resorufin (set as 100 %). Data are representative of three biological replicates; statistical significance was assessed using one-way ANOVA, followed by Tukey HSD *post-hoc* test, where letters indicate statistical differences between groups ($p$-value $< 0.05$).

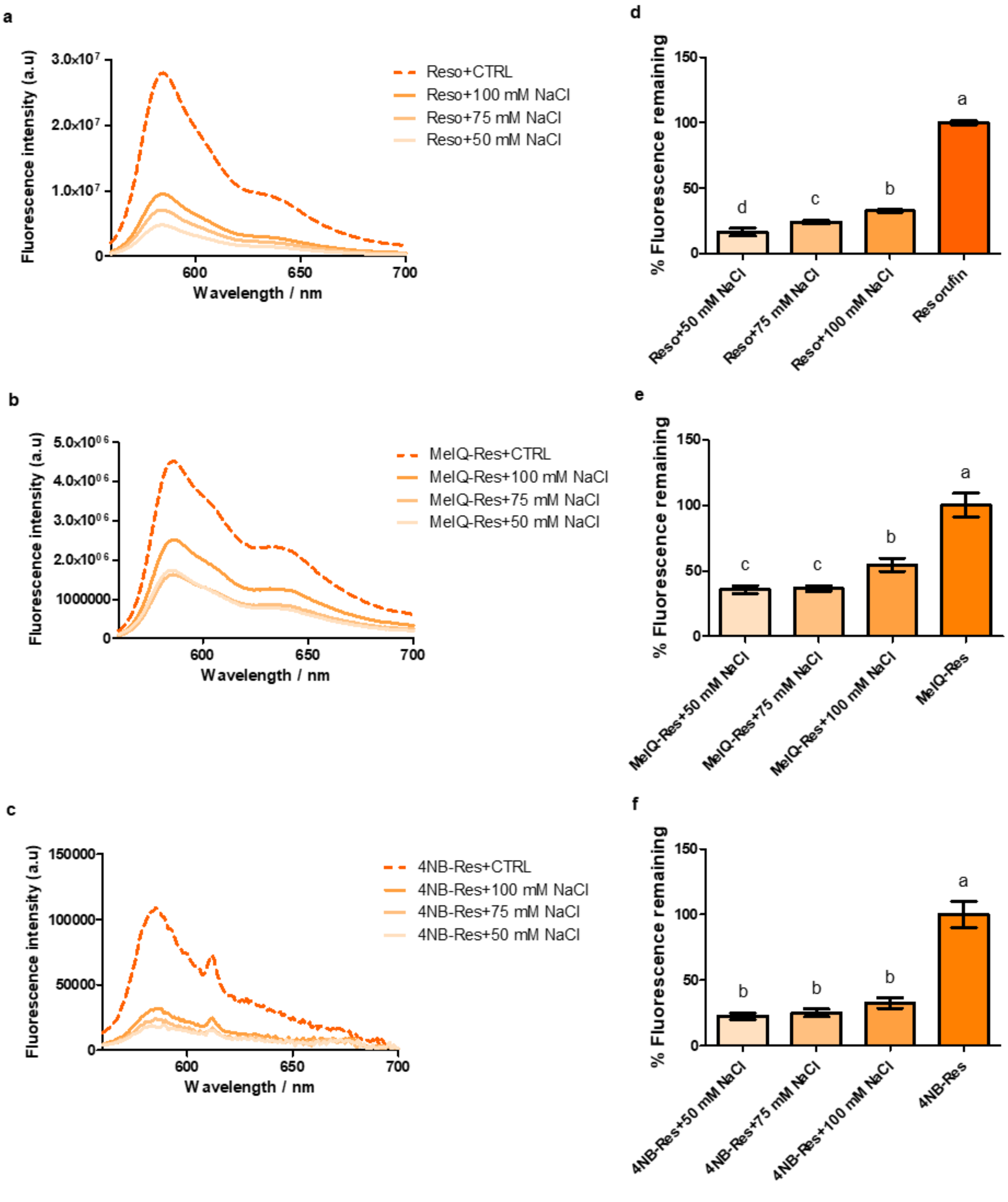
a
Fluorescence intensity (a.u)
3.0x10^7
2.0x10^7
1.0x10^7
0
600
650
700
Wavelength / nm
Reso+CTRL
Reso+100 mM NaCl
Reso+75 mM NaCl
Reso+50 mM NaCl
b
Fluorescence intensity (a.u)
5.0x10^06
4.0x10^06
3.0x10^06
2.0x10^06
1000000
0
600
650
700
Wavelength / nm
MeIQ-Res+CTRL
MeIQ-Res+100 mM NaCl
MeIQ-Res+75 mM NaCl
MeIQ-Res+50 mM NaCl
c
Fluorescence intensity (a.u)
150000
100000
50000
0
600
650
700
Wavelength / nm
4NB-Res+CTRL
4NB-Res+100 mM NaCl
4NB-Res+75 mM NaCl
4NB-Res+50 mM NaCl
d
% Fluorescence remaining
150
100
50
0
d
c
b
a
Reso+50 mM NaCl
Reso+75 mM NaCl
Reso+100 mM NaCl
Resorufin
e
% Fluorescence remaining
150
100
50
0
c
c
b
a
MeIQ-Res+50 mM NaCl
MeIQ-Res+75 mM NaCl
MeIQ-Res+100 mM NaCl
MeIQ-Res
f
% Fluorescence remaining
150
100
50
0
b
b
b
a
4NB-Res+50 mM NaCl
4NB-Res+75 mM NaCl
4NB-Res+100 mM NaCl
4NB-Res

### Chemistry experimental section

**Chemicals** were purchased from Sigma Aldrich UK, Alfa Aesar UK, and Fluorochem, and were used as supplied unless stated. Brine refers to a saturated aqueous solution of sodium chloride. Anhydrous solvents were obtained from an MBRAUN Solvent Purification System 5 and stored under an argon atmosphere over 3 Å molecular sieves. Petroleum ether refers to the fraction of light petroleum ether boiling in the range 40–60 °C. *In vacuo* refers to the removal of solvent using a Buchi® rotary evaporator under reduced pressure in a water bath at 40 °C.

**$^{1}$H NMR** spectra were recorded on Bruker AVC500 (500 MHz) or Bruker AVH400 (400 MHz) spectrometers using deuterochloroform (unless indicated otherwise) as a reference for the internal deuterium lock. The chemical shift data for each signal are given as $\delta_H$ in units of parts per million (ppm) relative to tetramethylsilane (TMS) where $\delta_H$ (TMS) = 0.00 ppm. The multiplicity of each signal is indicated by s (singlet); d (doublet); dd (doublet of doublets); or m (multiplet). The number of protons (n) for a given resonance signal is indicated by nH. Coupling constants (*J*) are quoted in Hz and are recorded to the nearest 0.1 Hz. Identical proton coupling constants (*J*) are averaged in each spectrum and reported to the nearest 0.1 Hz. The coupling constants are determined by analysis using MestreNova software. Bruker Topspin was used to plot the spectra. Spectra were assigned using COSY, NOESY, HSQC, and HMBC experiments as necessary.

**Mass spectra** were acquired on a VG platform spectrometer and an Agilent 6120 spectrometer (low resolution). Electro-spray ionization spectra were obtained on Micromass LCT Premier and Bruker MicroTOF spectrometers, operating in a positive or negative mode, as indicated, from solutions of MeOH or MeCN. *m/z* values are reported in Daltons and followed by their percentage abundance in parentheses. Electron ionization/field ionization (EI/FI) was carried out on a Waters GCT with a temperature-programmed solids probe inlet. Samples were introduced in glass tips directly into the source where they were vaporized and analyzed. In EI the ionization is by electron impact, electrons being provided by a filament. In FI the ionization is in an intense electric field which causes quantum electron tunneling of a valence electron.

**Melting points** were determined using a Griffin capillary tube melting point apparatus and are uncorrected.

**Analytical thin-layer chromatography (TLC)** was carried out on normal phase Merck silica gel 60 F254 aluminum-supported chromatography sheets. Visualization was done by absorption of UV light ($\lambda_{max}$ 254 and 365 nm) or thermal development after staining in an aqueous solution of potassium permanganate. UV light was provided by a LF – 206.LS 230V – 50 Hz from UVltec Limited

**Flash column chromatography** was performed manually using Geduran® silica gel 60 (40-63 µm) eluting with solvents as supplied, under a positive pressure of air or nitrogen on a Biotage® Selekt Flash Pure Purification System using Biotage® KP-Sil SNAP or Biotage® Sfär Silica cartridges.

***General procedures for reactive oxygen species (ROS) preparation and assays (Zhang et al., 2015)***

*General procedure for hydrogen peroxide ($H_2O_2$) assay:*

*Preparation*

The concentration of $H_2O_2$ (stock solution) was determined by measuring its absorption at 240 nm, using an extinction coefficient of 43.6 $cm^{-1}M^{-1}$. To prepare the assay solution, 15 µL of $H_2O_2$ (30 wt. % stock solution) was added to 5 mL of Milli-Q water in a 15 mL Falcon tube. Using the aforementioned method, the concentration of the resulting solution was determined to be 28.8 mM.

*Assay*

To a 3 mL quartz cuvette on ice (0–4 °C) containing Milli-Q water (pH 7.4, 2986.5 µL) and $H_2O_2$ solution (10.5 µL of a 28.8 mM stock solution in Milli-Q water), a probe solution (3 µL of a 1 mM stock solution in DMSO) was added. The cuvette was sealed and left at room temperature for 5 minutes. At this point, the final concentrations of $H_2O_2$ and the hypoxia-imaging probe were 100 µM and 1 µM, respectively. The fluorescence spectra were then measured using the specified parameters at the given time point.

*General procedure for hydroxyl radical (HO•) assay:*

*Preparation*

The hydroxyl radical (HO•) was generated using the Fenton reaction.To prepare the assay solution, 15 µL of $H_2O_2$ (30 wt. % stock solution) as added to 5 mL of Milli-Q water in a 15 mL Falcon tube, resulting in a 28.8 mM solution. From this, 1 mL of the $H_2O_2$ solution (28.8 mM, 10 eq.) was transferred to another 15 mL Falcon tube, and ferrous chloride (2.9 mM, 1 eq.) was added. The solution turned orange and was used immediately for the assay.

*Assay*

To a 3 mL quartz cuvette on ice (0–4 °C) containing Milli-Q water (pH 7.4, 2986.5 µL) and HO• solution (10.5 µL of a 28.8 mM stock solution in Milli-Q water), a probe solution (3 µL of a 1 mM stock solution in

DMSO) was added. At this stage, the final concentrations of HO• and the hypoxia-imaging probe were 100 µM and 1 µM, respectively. The solution was homogenized by pipetting for 10 s, then the cuvette was sealed and left at room temperature for 5 minutes. The fluorescence spectra were measured under the specified parameters.

***General procedure for light stability assay:***

To a 1 mL transparent vial containing DMSO (45 µL), a probe solution (5 µL of a 10 mM stock solution in DMSO) was added. An aliquot (3 µL) was taken from the reaction solution and transferred to a 3 mL quartz cuvette, followed by the addition of Milli-Q water (pH 7, 2997 µL). The solution was homogenized by pipetting for 10 s, then the fluorescence spectra were recorded using the specified parameters, serving as the initial time point (T = 0, where T represents time).The vial was then sealed and placed in an aluminum box, positioned 20 cm away from a white LED (OSRAM, 11 W, 52 mA, 1521 lm), and illuminated for 4 hours. After illumination, another aliquot (3 µL) was collected, transferred to a 3 mL quartz cuvette, and diluted with Milli-Q water (pH 7, 2997 µL). The fluorescence spectrum was subsequently recorded under the specified parameters.

***General procedure for the assay with high ionic strength buffers:***

To a 3 mL quartz cuvette containing buffer with the desired NaCl concentration (2997 µL), a probe solution (3 µL of a 1 mM stock solution in DMSO) was added. The solution was homogenized by pipetting for 10 s, then the cuvette was sealed and left at room temperature for 10 minutes. The fluorescence spectra were then recorded using the specified parameters.

General procedure for the assay with different pH buffers:

To a 3 mL quartz cuvette containing buffer at the desired pH (2997 µL), a probe solution (3 µL of a 1 mM stock solution in DMSO) was added. The solution was homogenized by pipetting for 10 s, then the cuvette was sealed and incubated at room temperature for 5 or 60 minutes. The fluorescence spectra were then recorded using the specified parameters.

**Synthetic procedures**

**7-[(4-Nitrobenzyl)oxy]-3H-phenoxazin-3-one (4-NB Resorufin)**

4-nitrobenzylbromide (89.1 mg, 0.412 mmol, 1.0 eq) in N,N-Dimethylformamide (1.5 mL) was added dropwise to a solution of resorufin (97.0 mg, 0.412 mmol, 1.0 eq) and potassium carbonate (85.5 mg, 0.618 mmol, 1.5 eq) in N,N-Dimethylformamide (3.5 mL). The solution was stirred at 40 ºC for 3 hours. The solution was diluted with ethyl acetate (100 mL) and washed with brine (3 × 50 mL). The aqueous layer was extracted with ethyl acetate (3 × 30 mL). The organic components were combined, dried (sodium sulfate), filtered and concentrated in vacuo. The crude residue was recrystallized from dichloromethane, followed by trituration with diethyl ether (3 × 10 mL) to yield the title compound (112 mg, 65%) as a red solid; $R_f$. 0.44 (50% ethyl acetate in petroleum ether), mp: decomposes >240 ºC (from diethyl ether), [Lit. (Collins *et al.*, 2018) 289–292 ºC]; $^1$H NMR (400 MHz, $CDCl_3$) δ 8.22 (2H, d, *J* = 8.6, *H-3’*), 7.68 (1H, d, *J* = 8.9, *H-9*), 7.56 (2H, d, *J* = 8.6, *H-2’*), 7.35 (1H, d, *J* = 9.7, *H-1*), 6.95 (1H, dd, *J* = 8.9, *J* = 2.7, *H-8*), 6.80 (1H, d, *J* = 2.7, *H-6*), 6.77 (1H, dd, *J* = 9.7, *J* = 2.0, *H-2*), 6.25 (1H, d, *J* = 2.0 , *H-4*), 5.22 (2H, s, C1’C$H_2$O).; LRMS m/z (ES+) 349.0 ([M+H]+, 4.27%). Analytical HPLC (Dionex Acclaim$^{TM}$ 120 C18 column [5 µm, 120 Å, 4.6 × 150 mm]; 95:5 $H_2O$: MeCN → 5:95 $H_2O$: MeCN: $H_2O$ with 0.1% TFA modifier, 10 min; 5 min hold; 1.5 mL min$^{-1}$] Ret. Time = 10.6 min. Purity: 98.4%. These data are in accordance with literature (Collins *et al.*, 2018).

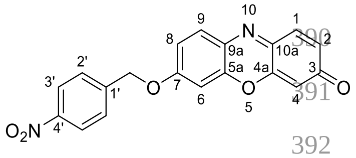


**5’-Methoxy-1’,2’-dimethyl-3-(1-((3-oxo-3*H*-phenoxazin-7-yl)oxy)ethyl)-1*H*-indole-4’,7’-dione (MeIQ Resorufin)**

Diisopropyl azodicarboxylate (0.120 mL, 0.602 mmol, 3.0 eq.) was added dropwise to a solution of methyl indolequinone (Wallabregue *et al.*, 2023) (50.0 mg, 0.201 mmol, 1.0 eq.), resorufin (107 mg, 0.502 mmol, 2.5 eq.) and triphenylphosphine (132 mg, 0.502 mmol, 2.5 eq.) in anhydrous tetrahydrofuran (3 mL) at room temperature under argon. The reaction mixture was stirred vigorously for 24 hours at 50 °C, then concentrated *in vacuo*. The residue was then dissolved in ethyl acetate (50 mL), washed with aqueous sodium hydrogen carbonate (3 × 20 mL), water (3 × 20 mL), aqueous hydrochloric acid (50 mL, 1 M solution), brine (2 × 20 mL), dried (sodium sulfate), and concentrated *in vacuo*. The crude material was purified using silica gel column chromatography eluting with petroleum ether:ethyl acetate (gradient 10 to 100% ethyl acetate). The residue was dissolved in ethyl acetate (2.5 mL) and precipitated by adding hexane (50 mL). The orange precipitate was filtered, washed with hexane (50 mL), and the precipitation procedure was repeated twice. The precipitate was then dried *in vacuo* to give the title compound as an orange solid (36 mg, 40%): $R_f$ 0.22 (petroleum ether:ethyl acetate, 1:1); m.p. 77–79 °C

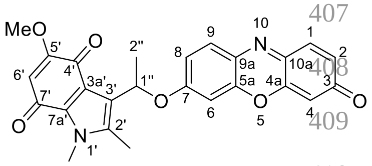

(ethyl acetate); [Lit. (Wallabregue *et al.*, 2023) 78 ºC]; $^{1}H$ NMR (600 MHz, $CDCl_3$) δ 7.62 (d, *J* = 8.9 Hz, 1H, *H-9*), 7.38 (d, *J* = 9.8 Hz, 1H, *H-1*), 6.92 (dd, *J* = 8.9, 2.6 Hz, 1H, *H-8*), 6.80 (dd, *J* = 9.8, 2.0 Hz, 1H, *H-2*), 6.78 (d, *J* = 2.6 Hz, 1H, *H-6*), 6.36 (q, *J* = 6.5 Hz, 1H, *H-1''*), 6.29 (d, *J* = 2.0 Hz, 1H, *H-4*), 5.66 (s, 1H, *H-6'*), 3.86 (s, 3H, $OCH_3$), 3.82 (s, 3H, $NCH_3$), 2.30 (s, 3H, (C-2')$CH_3$)), 1.68 (d, *J* = 6.5 Hz, 3H, *H-2''*); LRMS *m/z* ($ESI^+$) 445 ($[M+H]^+$, 90%); Analytical HPLC (Dionex Acclaim™ 120 C18 column [5 µm, 120 Å, 4.6 × 150 mm]; 95:5 $H_2O$: MeCN → 5:95 $H_2O$: MeCN: $H_2O$ with 0.1% TFA modifier, 10 min; 5 min hold; 1.5 mL $min^{-1}$] Ret. Time = 10.6 min. Purity: 93.5%. The spectroscopic data are in good agreement with literature values (Wallabregue *et al.*, 2023).

### HPLC Traces

**75’-Methoxy-1’,2’-dimethyl-3-(1-((3-oxo-3*H*-phenoxazin-7-yl)oxy)ethyl)-1*H*-indole-4’,7’-dione (MeIQ Resorufin) @254 nm**

**OXAW51-2** **10/2/2021**

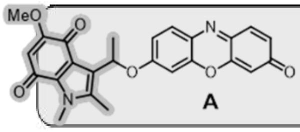


**A**

| | |
|---|---|
| Acquisition Method | Purity short run @254 nm |
| Acquisition Date/Time | 9/13/2021 4:46 pm |
| Injection Volume | 40 |
| Sample Name | OXAW51-2 |
| Sample Description | |
| Batch Description | |

OXAW51-2 : Injection 1

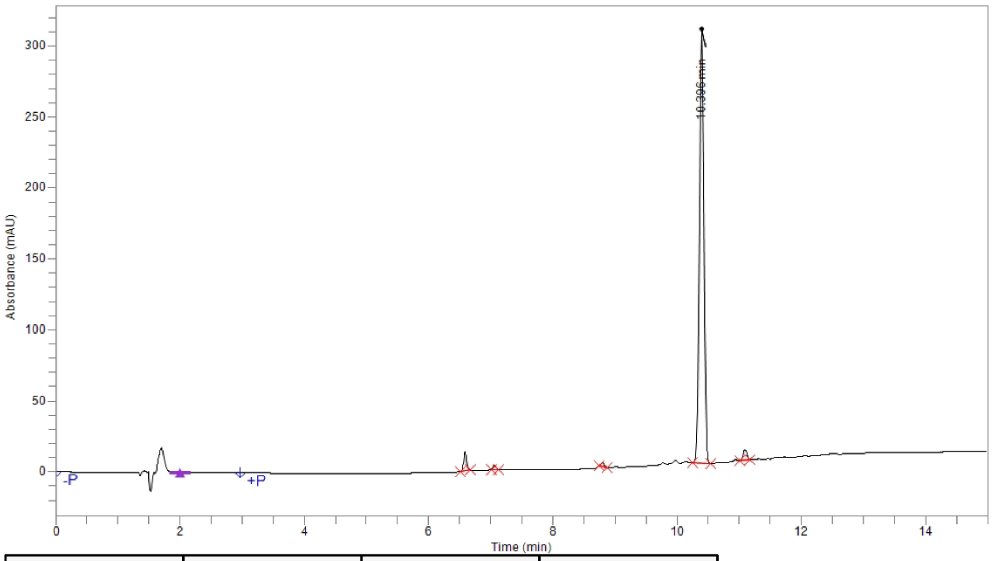


| Time | Height | Area | Area % |
|---|---|---|---|
| 6.588 | 13,446.3 | 47,468.8 | 3.13 |
| 7.065 | 2,714.3 | 8,841.8 | 0.58 |
| 8.798 | 2,791.6 | 11,067.8 | 0.73 |
| 10.396 | 305,851.7 | 1,417,074.4 | 93.53 |
| 11.093 | 7,270.8 | 30,670.2 | 2.02 |
| **Total** | | 1,515,123.0 | 100.00 |

**7-[(4-Nitrobenzyl)oxy]-3H-phenoxazin-3-one (4-NB Resorufin) @254 nm**

**OX-AW91@254** **6/4/2021**

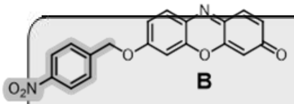


**B**

| | |
|---|---|
| Acquisition Method | Purity short run @254 nm |
| Acquisition Date/Time | 6/3/2021 6:10 pm |
| Injection Volume | 30 |
| Sample Name | OX-AW91@254 |
| Sample Description | |
| Batch Description | |

OX-AW91@254 : Injection 1

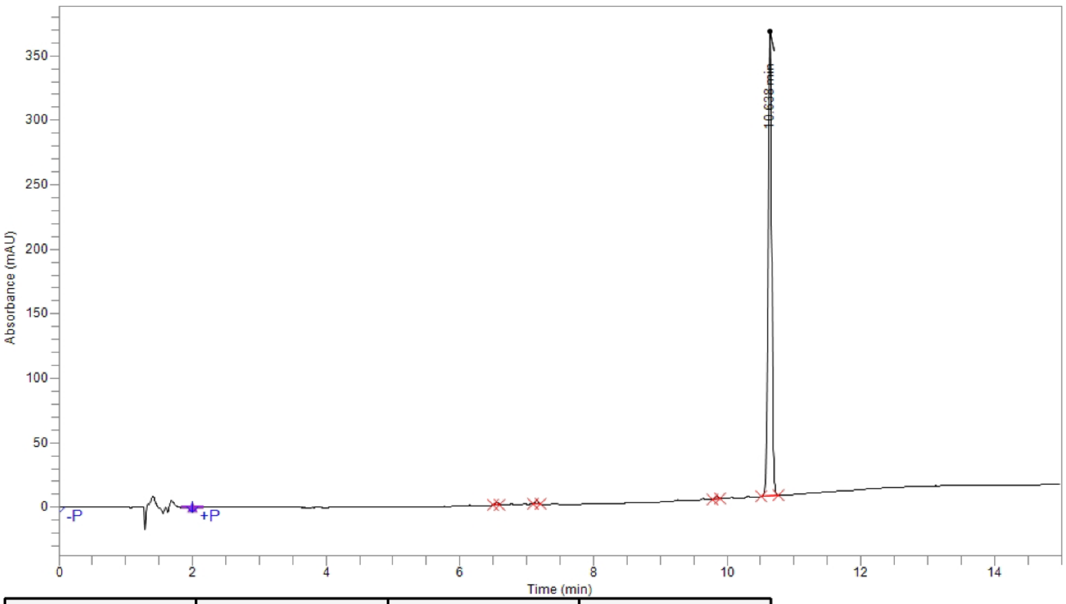


| Time | Height | Area | Area % |
|---|---|---|---|
| 6.550 | 1,914.1 | 6,323.4 | 0.47 |
| 7.148 | 3,003.3 | 8,985.0 | 0.67 |
| 9.844 | 2,022.7 | 6,576.5 | 0.49 |
| 10.638 | 360,908.8 | 1,318,842.1 | 98.37 |
| **Total** | | 1,340,727.0 | 100.00 |
